# Dynamic heterogeneity in an *E. coli* stress response regulon mediates gene activation and antimicrobial peptide tolerance

**DOI:** 10.1101/2024.11.27.625634

**Authors:** Caroline M. Blassick, Hossein Moghimianavval, Fereshteh Jafarbeglou, Jean-Baptiste Lugagne, Mary J. Dunlop

## Abstract

The bacterial stress response is an intricately regulated system that plays a critical role in cellular resistance to drug treatment. The complexity of this response is further complicated by cell-to-cell heterogeneity in the expression of bacterial stress response genes. These genes are often organized into networks comprising one or more transcriptional regulators that control expression of a suite of downstream genes. While the expression heterogeneity of many of these upstream regulators has been characterized, the way in which this variability affects the larger downstream stress response remains hard to predict, prompting two key questions. First, how does heterogeneity and expression noise in stress response regulators propagate to the diverse downstream genes in their regulons? Second, when expression levels vary, how do upstream and downstream genes act together to protect cells from stress? To address these questions, we focus on the transcription factor PhoP, a critical virulence regulator which coordinates pathogenicity in several gram-negative species. We use optogenetic stimulation to precisely control PhoP expression levels and utilize information theory to examine how variations in PhoP affect the downstream activation of genes in the PhoP regulon. We find that these downstream genes exhibit differences in mean expression level, sensitivity to increasing levels of PhoP, and signal transmission reliability. These response functions can also vary between individual cells, increasing heterogeneity in the population. We tie these variations to cell survival when bacteria are exposed to a clinically-relevant antimicrobial peptide, showing that while high expression of the PhoP-regulon gene *pmrD* provides a protective effect against Polymyxin B, cell survival chances are best determined by integrated dynamic PhoP and *pmrD* expression levels. Overall, we demonstrate that cell-to-cell and temporal heterogeneous expression of a stress response regulator can have clear consequences for enabling bacteria to survive stress.

## Introduction

Bacteria rely on complex, interconnected gene regulatory networks to maintain homeostasis and tolerate a diverse range of external stressors, including osmotic stress, heat or cold shock, nutrient starvation, acidic environments, and treatment with antimicrobial agents.^1,2^ These environmental responses are often initiated by two-component systems, a set of critical regulators responsible for mediating responses to these stressors and triggers.^3–5^ Receptors sense a stressor through direct ligand binding or other mechanisms, and the receptor’s histidine kinase phosphorylates its response regulator, which then serves as a transcription factor to modulate downstream gene expression. Gene regulatory networks may demonstrate considerable complexity through multi-layer cascades,^6^ combinatorial regulation,^7^ or feedback control.^8^ While extensive progress has been made in modeling the behavior of these regulatory networks,^9,10^ the precise relationship between the dynamics of a particular transcription factor and how it regulates genes within the stress response network is not always straightforward to determine.^11,12^

Additionally, transcription factors and other genes in stress response pathways can be heterogeneously expressed in isogenic populations, even in the absence of external stressors.^13,14^ Heterogeneous gene expression stems from both intrinsic and extrinsic events, including transcriptional bursting dynamics, stochastic partitioning of molecules during cell division, competition over limited resources, and fluctuating environmental factors.^13,15,16^ Stochastic, heterogeneous gene expression can have important functional consequences, for instance by enabling “bet-hedging” strategies that allow subsets of cells to survive stress.^17,18^ This protective effect has also been implicated in the development of antibiotic resistance, by providing windows of tolerance in which cells have time to evolve more permanent resistance mechanisms.^19^ Fully characterizing stress response networks therefore requires understanding of not only average gene expression patterns, but also the dynamics and extent of cell-to-cell heterogeneity and how this heterogeneity propagates through regulatory networks to influence stress tolerance.

The PhoPQ stress response network has been well-studied due to its role in virulence in *Escherichia coli* and other gram-negative species, such as *Salmonella enterica* serovar Typhimurium and *Pseudomonas aeruginosa.*^20,21^ PhoPQ is a two-component system consisting of the membrane-associated sensor PhoQ and its response regulator, PhoP. PhoQ can either phosphorylate or de-phosphorylate PhoP depending on environmental conditions, including low Mg²⁺ or the presence of cationic antimicrobial peptides.^22^ When phosphorylated, PhoP activates genes involved in Mg²⁺ transport, outer membrane remodeling, acid resistance, and antimicrobial peptide tolerance.^23^ For example, expression of *pmrD* has been implicated in tolerance of Polymyxin antimicrobial peptides.^24,25^ While a relatively small number of promoters contain motifs annotated as PhoP binding boxes, deletion of PhoP results in the differential expression of over 600 genes.^26^ The diverse and far-reaching effects of PhoP activation motivated our focus on the PhoP regulon in order to understand the role of differential regulation and heterogeneity across a stress response network.

Prior studies comparing measurements obtained from gel shift assays that were focused on promoters directly bound by PhoP have found that the affinity of PhoP for each of these promoters can differ.^23^ Intriguingly, PhoP overexpression data *in vivo* can qualitatively disagree with these *in vitro* measurements. In one study, fluorescent reporters were used to track expression from several genes in the PhoP regulon over a range of stimulus levels. While the fold change in expression of each gene was similar with each change in Mg^2+^ concentration, there were large differences in the baseline expression level of each gene.^27^ As levels of phosphorylated PhoP increase even further in cells, however, differential regulation patterns may emerge, with different genes exhibiting different fold changes or saturation points in response to the same stimulus.^27^ These variable characteristics can help explain the differential regulation of genes that are all controlled by the same transcription factor. However, transcription factor binding affinity and saturation points alone are not the sole determinants of downstream response, thus a more comprehensive understanding of this pathway requires analysis of rich expression data in living cells.^28^

The input-processing network in a cell is made up of signaling molecules that are known to act as both sensors of cell state and functional regulators of cell response when facing changes in the cell environment. Therefore, while the regulatory network of genes in the cell consists of many input-output relationships, the network itself acts as a unit to perceive the cell state and drive functionally relevant outputs.^29–32^ Traditional approaches for characterizing gene networks, such as the construction of gene knock-out strains,^33^ inherently fail to capture the rich temporal dynamics that underlie gene network regulation.^34^ Dynamic perturbations, such as varying the amount of a given transcription factor over time, can capture more information than a single snapshot, enabling more accurate prediction of cell fate.^31,35,36^ Additionally, capturing input-output dynamics allows for the temporal characterization of cell-to-cell heterogeneity or noise in gene expression.^37^

Optogenetic, or light-based, regulation of gene expression provides a powerful means of dynamically perturbing cells with precise spatial and temporal control.^38^ Additionally, optogenetic regulation can be linked with hardware controllers for feedback-enabled regulation of the gene of interest.^39^ This makes optogenetic control an especially attractive option for obtaining rich, single-cell datasets with high reproducibility.

To examine the dynamics and cell-to-cell heterogeneity of the PhoP regulon, we leveraged the advantages of optogenetic control to modulate levels of PhoP. We then traced the effect of PhoP activity level to a suite of downstream genes using fluorescent reporters. In both bulk culture and at the single-cell level, we found marked differences in the activation patterns of different reporters, as quantified by mutual information analysis.

In addition to end-point bulk culture measurements, we used microfluidics and high-throughput microscopy to capture dynamic heterogeneous gene expression at the single-cell level. Precise optogenetic control and single-cell gene expression data also revealed new insights into the role that cell-to-cell heterogeneity plays in this important stress response network. Within each individual gene, we found broad consistency in the average response of cells to different levels of stimuli. However, we uncovered distinct, gene-specific, and heterogeneous dynamic input-output distributions. The effects of these differences in dynamic input and output signals, which we present as joint time series data, enable interpretation of meaningful differences in how regulatory pathways encode information. Using these heterogeneous dynamic signals, we analyzed cells after treatment with the antimicrobial peptide Polymyxin B, finding that these joint times series were predictive of cell fate. In addition, we found that while the PhoP-controlled gene *pmrD* plays a role in resistance, high levels of *pmrD* alone are not strong determinants of survival. Rather, it is the dynamic joint PhoP/*pmrD* space that best predicts survival, meaning that an interplay of both upstream and downstream regulators determines the probability of survival.

Altogether, our results demonstrate that variable input-output relationships between a stress response transcription factor and its regulon play an important role in shaping the dynamics and functional consequences of network activation. Furthermore, we showcase a powerful platform with which the dynamics and heterogeneity of regulatory pathways can be investigated at the single-cell level.

## Results

### Optogenetic system for control of gene activation

To enable rapid and precise control, we used an optogenetic platform for activation of PhoP. For real-time tracking of downstream network dynamics, we used PhoP-regulated gene reporters to examine the effect of PhoP perturbation on a diverse subset of its regulon. While there is evidence that wild-type PhoP can bind the promoters of downstream genes in an unphosphorylated (*i.e.*, not activated) state,^40^ it is most active when phosphorylated by the sensor kinase PhoQ under low Mg^2+^ conditions. PhoP then activates transcription of a set of genes with diverse functions (Fig. 1A). To select PhoP regulon genes for investigation, we first narrowed down to a subset of genes with strong evidence of direct control via PhoP and further narrowed the subset by removing genes with low baseline expression as well as low input sensitivity, to ensure that expression noise could be reliably measured. We chose a final subset from these candidates to represent a wide array of functions. Specifically, we selected *hemL*, a glutamate-1-semialdehyde aminotransferase involved in porphyrin biosynthesis;^41^ *nagA*, an N-acetylglucosamine-6-phosphate deacetylase involved in peptidoglycan recycling;^42^ *pmrD*, a homolog to a protein implicated in Polymyxin B resistance in *Salmonella enterica*;^43^ *rstA*, a transcriptional regulator associated with virulence in pathogenic strains of *E. coli*;^44,45^ *slyB*, a membrane lipoprotein important for maintaining membrane integrity during stress;^46^ and *yrbL*, an uncharacterized protein previously found to be regulated by PhoP.^23^ PhoP can additionally activate transcription from its own promoter in the form of positive autoregulation, thus we also included *phoP*.^20^ Fig. S1 provides details on PhoP regulation and other regulatory factors for the selected genes.

**Figure 1.**
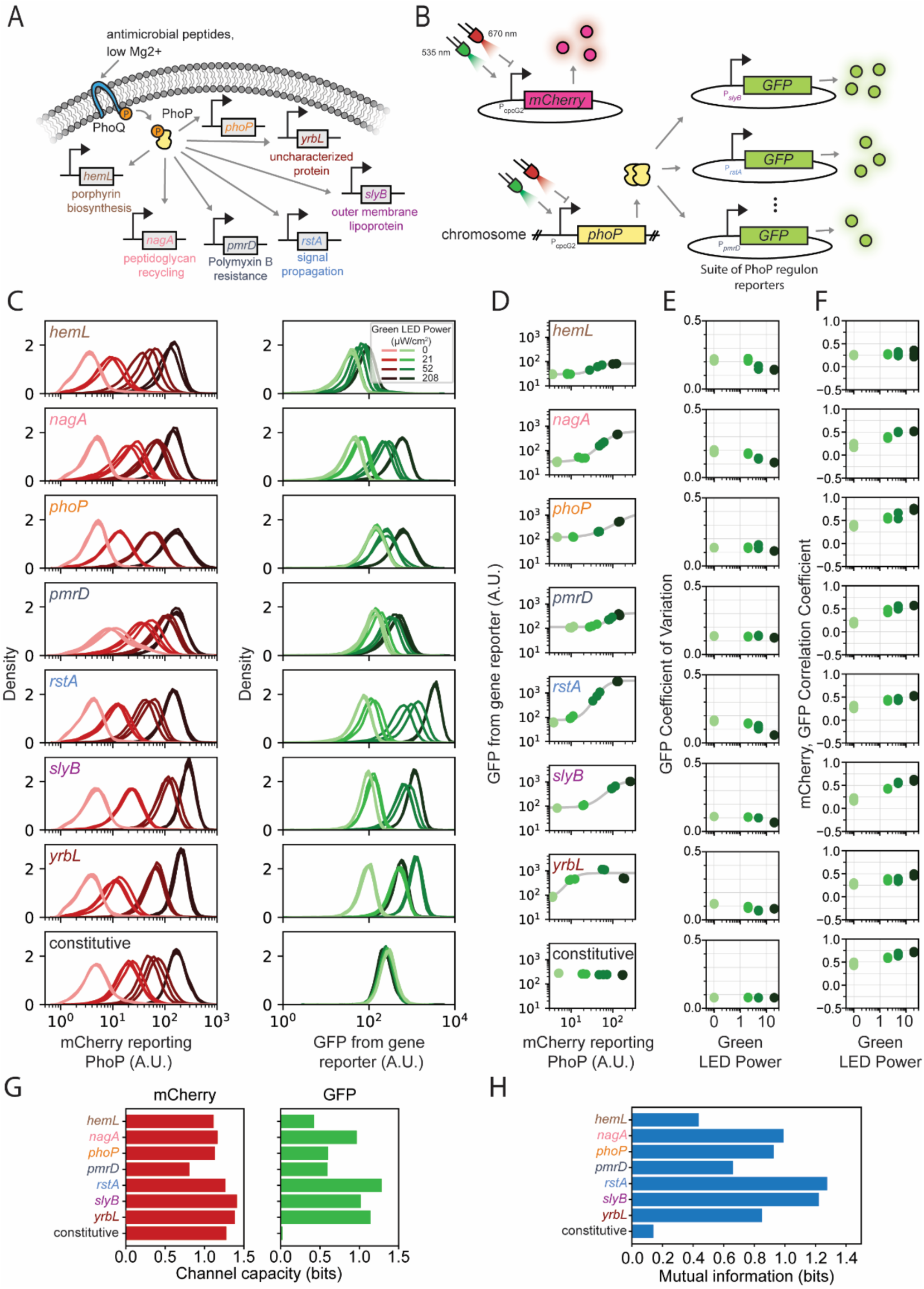
Optogenetic control of PhoP in bulk culture. (A) Schematic of the PhoPQ two-component system. Upon stimulation by low Mg^2+^ or the presence of antimicrobial peptides, there is a net increase in the rate of PhoP phosphorylation by the sensor kinase PhoQ. Phosphorylated PhoP activates transcription from a wide range of genes, a subset of which are included here. (B) Optogenetic regulation of PhoP. *phoP* is placed downstream of the optogenetic P*_cpcG2_* promoter along with mCherry. Increasing levels of green light result in increased production of PhoP, which then activates transcription of *gfp* from PhoP-regulon reporter plasmids. (C) mCherry and GFP fluorescence distributions for cells containing the system described in panel B, under a range of green light stimulation levels. n = 3 technical replicates for each green light level. See Fig. S15 for biological replicates. (D) Geometric means of GFP and mCherry fluorescence of the cells from panel C. Grey line represents best fit to a Hill function model. (E) Coefficient of variation for GFP across all cells. See Table S2 for statistical significance between the groups. (F) Mean correlation coefficient between mCherry and GFP calculated across all individual cells. See Table S3 for statistical significance between the groups. (G) Channel capacity of mCherry (left) or GFP (right) populations given the four optogenetic stimulation levels. (H) Mutual information between continuous mCherry (PhoP) and GFP (downstream gene) distributions.

The low Mg^2+^ conditions that result in maximal phosphorylation of PhoP can be severely growth limiting (Fig. S2), which complicates culturing and controlling cells. To circumvent this, we employed the strategy of Miyashiro *et al*. by using a mutated version of PhoP, which behaves as though constitutively phosphorylated.^27^ Unless otherwise noted, it is this mutated version of PhoP that is used for all optogenetic experiments.

To enable precise control of PhoP dynamics, we employed the CcaS/R two-component optogenetic system.^47^ In this system, the CcaS sensor kinase enters its active conformation upon exposure to green light (535 nm), allowing it to phosphorylate the CcaR response regulator. Phosphorylated CcaR then activates transcription from the optogenetic P*_cpcG2_* promoter. Red light (670 nm) reverses this process, repressing transcription. We integrated the PhoP transcription factor under the control of the P*_cpcG2_* promoter into the chromosome of BW25113 Δ*phoP* cells, allowing its levels to be increased when cells were exposed to green light and decreased when cells were exposed to red light (Fig. 1B). We used the Δ*phoP* strain to prevent confounding effects from variation in native *phoP* expression. To have a readout of transcription factor production, we also placed the gene encoding the red fluorescent protein mCherry under control of the P*_cpcG2_* promoter.

As a measure of downstream activation, we utilized a set of reporter plasmids,^48^ which consists of the gene encoding green fluorescent protein GFP under the control of a promoter known to be regulated by PhoP. To confirm that CcaS/R activation does not have spurious effects on cell transcription, such as global transcriptional regulation that is independent of PhoP-mediated regulation, we also included a constitutive promoter reporter as a negative control. This dual fluorophore setup allowed signal propagation to be traced from the PhoP transcription factor to downstream genes by comparing the levels of mCherry and GFP. We note that although mCherry and GFP have different maturation dynamics compared to each other^49^ as well as to the proteins they report, since our measurements in all experiments are performed with identical reporters, it is possible to compare their relative differences in expression patterns.

Using a suite of different GFP reporters, each under the promoter of a different gene driving GFP expression, reveals how identical upstream transcription factor dynamics can result in different expression patterns from downstream genes. Additionally, comparing individual cells containing identical reporters can provide insights into cell-to-cell heterogeneity in gene expression and its propagation.

### Optogenetic control of PhoP

We began by optogenetically stimulating the PhoP light-controlled strains and measuring the corresponding change in fluorescence from each gene reporter (Fig. 1C). Using an optogenetic well plate equipped with red and green LEDs whose intensities can be independently controlled, we subjected cells containing the optogenetic PhoP system to a range of green light intensities.^50^ After around five hours of light exposure, which is sufficient for cells to reach steady-state expression levels under similar conditions,^51^ we used flow cytometry to measure mCherry and GFP fluorescence for all cells. mCherry levels, which report the amount of PhoP induction, increased with increasing levels of green light stimulation, as expected, and showed similar profiles across all experiments. In addition, we observed an overall increasing pattern for GFP as the optogenetic stimulation levels increased, though the details of the increase were highly dependent on the specific downstream gene.

To ensure that mCherry and GFP fluorescence measurements correspond to expression levels of PhoP and the downstream genes, respectively, we quantified the mRNA levels of *phoP* and *mCherry* as well as *slyB* (as a model downstream gene) and *gfp*. Our qPCR measurements showed a consistent monotonic increase in both *phoP* and *mCherry* expression levels with increasing green light intensity (Fig. S3A), confirming that *mCherry* levels are proportional to *phoP* levels. Similarly, both *slyB* and *gfp* expression increased with higher green light stimulation (Fig. S3B). We also checked whether physiological expression levels of *phoP* under normal and low Mg^2+^ conditions are comparable to light-induced *phoP* expression levels. qPCR measurements demonstrated low *phoP* expression when wild-type BW25113 cells were grown in lysogeny broth (LB) media and elevated *phoP* expression when cells were grown in low Mg^2+^ media (Fig. S3C).

Importantly, the range of *phoP* expression levels achieved via optogenetic stimulation included the physiological *phoP* expression level (Fig. S3A and C), indicating that the optogenetic system operates within a suitable dynamic range. Collectively, our qPCR measurements show *phoP* levels expressed via optogenetics are around physiological *phoP* levels, and that *mCherry* and *gfp* are appropriate proxies for *phoP* and downstream gene abundance.

Next, we set out to characterize the differences between downstream gene expression patterns in response to identical upstream variations. We observed that while all reporters for downstream genes showed an increase in GFP as optogenetic stimulation levels (*i.e.*, PhoP levels) increased, there were notable differences in the response patterns, as well as in the baseline expression levels of each gene. Some genes, such as *rstA*, reliably demonstrated a wide range of increases commensurate with all levels of optogenetic stimulation; others, such as *hemL*, only exhibited slight expression increases with higher levels of stimulation. These differences in activation dynamics result in differently shaped response curves when GFP levels are compared as a function of mCherry levels (Fig. 1D). Fits to a Hill function model confirm the wide diversity of response profiles that these downstream genes exhibit in response to identical PhoP signals (Fig. 1D, Table S1). As expected, the constitutive control exhibited minimal response to increasing optogenetic stimulation.

The distinct distributions of mCherry and GFP populations across different genes reflect the heterogeneous nature of gene expression. While the average expression follows a unique pattern for each gene (Fig. 1D), the response to elevated PhoP is different in each cell, as revealed by the distribution characteristics of the GFP signal. Because this heterogeneity in gene expression is especially important for stress response,^14,52^ we next examined variations in expression level within single cells using the flow cytometry data. Populations with similar mean fluorescence levels can have differing population distributions, for instance, exhibiting a wide distribution of responses to the same input. There may also be differences in the correlation between mCherry and GFP values for individual cells, demonstrating a difference in how reliably the target gene responds to a change in PhoP expression (Fig. S4). We began by asking whether the width of the distribution of gene expression changed with increasing stimulation levels, since noise in gene expression is known to decrease as the copy number of its regulators increases.^15^ We quantified this using the coefficient of variation. Despite differences in expression patterns, each reporter strain had broadly similar coefficients of variation for GFP across all stimulation levels (Fig. 1E, see Table S2 for statistical significance values). As an additional check, we also calculated Jensen-Shannon divergence, an unbiased metric for quantifying differences between two distributions. The Jensen-Shannon divergence among fluorescence measurements for each gene was close to zero (Fig. S5), indicating little difference between the width and shape of upstream vs. downstream fluorescence distributions within each individual gene. These results suggests that at the levels of PhoP activation explored in this study, there is not a large difference in the noise in downstream gene expression between the lowest and highest levels of PhoP activation. We note that since fluorescent protein reporters of PhoP and downstream gene––mCherry and GFP, respectively––are both encoded on plasmids, the extrinsic noise from variable plasmid copy numbers could, in principle, mask subtle changes in regulator or downstream gene copy number. However, because all measurements are subject to these same limitations, we can still draw conclusions about signal transmission properties based on differences between reporter outputs given identical inputs.

Next, we calculated the correlation coefficient between mCherry and GFP values for each individual cell. This metric indicates how tightly correlated the two values are, providing an indication, at the single-cell level, of how sensitive the downstream gene is to changes in the upstream regulator. We observed a general trend of increasing downstream sensitivity for most genes (Fig. 1F). This suggests that within single cells, downstream genes generally covary positively with PhoP changes, although with varying sensitivities.

We were next interested in quantifying the reliability of signal transduction from input (PhoP) to output (downstream gene). Although mCherry distributions follow similar trends for different downstream genes, the GFP populations have varying degrees of overlap with each other for each gene, meaning that a wide range of inputs can cause similar or distinct outputs. Unreliable, noisy signal transmission leads to overlapping outputs that are difficult to discriminate, while high-fidelity signal transmission leads to distinct populations. Drawing inspiration from information theory, signal transmission fidelity in various biological pathways has been characterized through mutual information.^31,53–56^ Mutual information, often reported in bits, implies the certainty of determining the input, knowing the output. Therefore, overlapping output distributions lead to low mutual information because determining their input, or the level of upstream regulator, from measured output is unreliable.

We first calculated the maximum mutual information, or channel capacity, of mCherry and GFP populations for each gene. Channel capacity determines the maximum amount of information that a discrete input channel can encode. Here, the different levels of optogenetic stimulation are considered input channels and the mCherry or GFP distributions are the corresponding outputs. Completely distinct and separate output distributions yield maximum channel capacity which is log_!_(# 𝑜𝑓 𝑖𝑛𝑝𝑢𝑡𝑠) bits. Here, given the 4 stimulation levels, this means that the maximum channel capacity is 2 bits, corresponding to perfect information transmission. We found that while mCherry channel capacity was generally around 1 bit, meaning there is enough information transmission for binary discrimination (on/off input), GFP channel capacity varied across a wide range for different genes (Fig. 1G). Genes such as *rstA* with low overlap between their GFP populations had a high channel capacity while *hemL* which showed less distinct GFP distributions had lower channel capacity. Expectedly, the constitutive reporter, being unresponsive to optogenetic stimulation, exhibited a GFP channel capacity close to zero.

Although channel capacity provides insight about how reliably PhoP or a downstream gene respond to optogenetic stimulations, a more accurate characterization of signal transmission fidelity in signaling pathways can be obtained by calculating mutual information between mCherry populations and GFP populations. Since each optogenetic input leads to a heterogeneous population of mCherry (PhoP) and GFP (downstream gene) expression, information theoretic analysis of the PhoP-downstream gene relationship as continuous variables can provide a more realistic input-output characterization of each pathway.^55^ Mutual information between mCherry and GFP distributions for each gene yielded generally similar results to channel capacity (Fig. 1H). However, for *slyB* and *phoP*, a higher mutual information was observed when continuous input-output relationships were taken into account. This indicates that the heterogeneity of input and output gene expression for *slyB* and *phoP* covaried, leading to more reliable information transmission. Conversely, the mutual information of *yrbL* was lower than its discrete-input channel capacity, meaning that variations in heterogeneous gene expression were independent for input and output, thus introducing noise to the signal transduction channel.

### Single-cell PhoP regulon dynamics

Dynamic phenotype switching or bet-hedging within heterogeneous populations of cells can be an important strategy to enable subsets of the population to withstand diverse types of stress. To explore whether this is the case in the PhoP network, we next asked whether the dynamics of PhoP regulation differ among the diverse genes in the regulon, and whether there is substantial heterogeneity both among individual cells and over time for single cells. In order to answer these questions, we grew a subset of the reporter strains in the mother machine microfluidic device to enable single-cell resolution measurements over many generations of cell growth.^57^ For these experiments, we focused on the reporters *phoP*, *pmrD*, *slyB*, and *yrbL*. We chose these reporters because they represent a diverse subset of the activation patterns observed in the bulk culture experiments, while also having similar baseline fluorescence levels, allowing them to be imaged under identical microscopy settings. We additionally included the constitutive reporter strain as a negative control.

Once cells were loaded into the mother machine, they were subjected to a five-minute loop that consisted of imaging in phase contrast, mCherry, and GFP channels, as well as optogenetic stimulation. Because the cells are trapped within individual chambers in the mother machine, we were able to use a digital micromirror device (DMD) to provide optogenetic stimulation separately to each individual chamber, allowing us to collect data for three objective groups within the same experiment (Fig. 2A). Each microscopy experiment began with four hours of red DMD stimulation applied to all cells to homogenize their expression levels. After hour four, cells were randomly divided into three subpopulations, each receiving a different optogenetic stimulation designed to drive mCherry (PhoP) expression toward a distinct objective: low, moderate, or high levels (Fig. 2B). After eight hours under these illumination conditions, all cells were returned to a low state with red DMD stimulation for the remainder of the experiment. To ensure that light stimulation during the experiment did not induce unintended growth effects, we compared the growth rate of cells grown in the dark with the cells grown under the three light exposure conditions and found that cells in all conditions grew comparably to cells maintained in the dark (Fig. S6).

**Figure 2.**
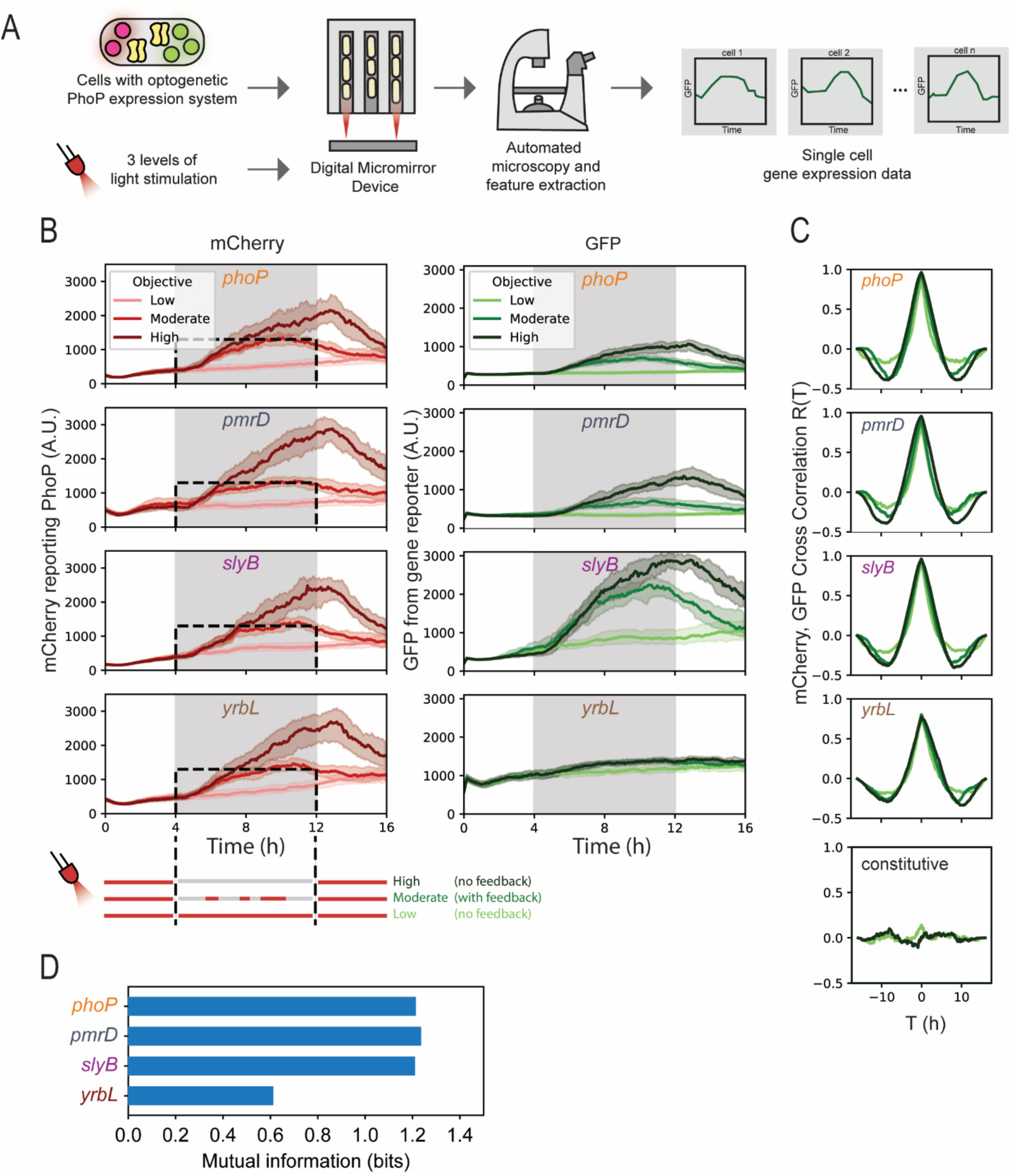
Single-cell optogenetic control of PhoP. (A) Experimental setup. Mother cells containing the PhoP optogenetic system are trapped in the end of the mother machine microfluidic device, allowing for long-term imaging. A Digital Micromirror Device provides individual stimulation to each mother machine chamber. Cells are imaged with automated microscopy and segmented on-the-fly, allowing for real-time monitoring of mCherry and GFP fluorescence. (B) mCherry and GFP expression levels over time for cells expressing optogenetically controlled PhoP. For the first and last four hours of each experiment, all cells receive red stimulation. Between hours 4 and 12, cells are randomly assigned to one of three objective groups: low, which receives constant red stimulations; high, which receives no red stimulations; and moderate, which is controlled to maintain 1300 A.U. mCherry fluorescence levels. Median plus error bars representing the middle quintile are shown for each objective group. n > 100 individual cells for each objective group. (C) Median values of cross-correlations computed between mCherry and GFP for every individual cell over time. Data for the constitutive control at low and high objective groups are also included. (D) Mutual information between multivariate dynamic mCherry (PhoP, input) and GFP (downstream gene, output) signals for each gene. Mutual information was calculated based on the dynamic mCherry and GFP signals only during the active control period, corresponding to hours 4 to 12 of the experiment.

In contrast to the flow cytometry data conditions, where we subjected bulk cultures to varying levels of green light illumination using a plate-based device and then measured mCherry and GFP fluorescence at a single timepoint, in microscopy experiments fluorescence imaging occurs every five minutes, thus providing time series of PhoP (mCherry) and downstream gene (GFP) variations. We found that the green light exposure from fluorescence imaging alone was sufficient to fully activate the CcaS/R optogenetic system, and that to achieve lower levels of expression we could use red light to repress expression. In the low stimulation subpopulation, cells were subjected to repressing red light from the DMD every five minutes. In the high stimulation subpopulation, cells received no DMD stimulation. However, as the data in Figure 2 show, the same level of light induction can produce variable levels of CcaS/R activation. Therefore, to achieve a similar moderate level of CcaS/R activation in all cells, we employed our previously described Deep Model Predictive Control platform^58^ (Fig. S7) to generate tailored DMD stimulation patterns for each cell, creating a constant level of CcaS/R activation and resulting in a constant mCherry objective of 1300 A.U. We note that in all three objective groups, the median expression follows an objective-specific pattern. However, due to stochastic and heterogeneous gene expression within each group, the exact value of mCherry or GFP is different among cells and fluctuates over time for a single cell.

This method of control allowed each reporter strain to have closely comparable mCherry levels over time, even when acquired in separate experiments (Fig. 2B). The median mCherry of the moderate stimulation population is very close to the set point for each experiment, and even without feedback control, the mCherry levels of the low and high objective populations show good agreement between experiments. The three induction levels also have distinct activation dynamics. The high objective population accumulates mCherry over time and shows continued expression past the 12 hour timepoint when repressive light is applied. Similarly, the low objective population slowly increases over time, likely due to the slight activation associated with illumination from image acquisition, including stimulation during fluorescence imaging. The control algorithm maintains the moderate objective, reached after five hours of controlled illumination, and mCherry also decays slowly after constant red light is applied to repress activation. These results demonstrate that we were able to reliably place cells at low, medium, and high levels of expression, allowing us to then investigate the more subtle effects of noise and heterogeneity within these populations.

All reporter strains also showed a corresponding increase in GFP activity upon stimulation of PhoP, with GFP levels rising the most for the *slyB* strain and the least for *yrbL* (Fig. 2B). As with the flow cytometry data, these results show clear differences in sensitivity to PhoP levels across the different gene reporters, despite all being controlled by the same transcription factor, revealing gene-specific signal transmission. Low sensitivity for some genes, such as *yrbL*, may be explained by the low saturation point that we also observed in the bulk culture experiments. While it was possible to conduct bulk culture experiments in the absence of all light except a repressing red wavelength, microscopy experiments required imaging with multiple other wavelengths that partially overlap with the activation spectrum of CcaS/R. This results in a small amount of activation of PhoP, and thus downstream genes expression, even under red DMD illumination and may result in less sensitivity for reporters, as we see in the case of *yrbL*. As a control, we verified that the GFP signal in the control strain carrying the light-inducible PhoP system along with a constitutive GFP reporter showed no light sensitivity (Fig. S8).

Since single-cell data enable analysis of dynamic signals, we set out to characterize the relationship between dynamic PhoP time series (input) on downstream gene expression (output). Computing the cross-correlations for each strain over time revealed that mCherry and GFP levels are highly correlated for all cells, achieving maximum values approaching 1 for all genes but *yrbL* (Fig. 2C). While equal increases in mCherry will not produce identical changes in GFP across the reporters, the high correlations that we observe indicate that for any deviation from the mean mCherry fluorescence, promoter activity from each reporter gene is very likely to deviate in the same direction. This relationship holds true across all objective groups. In addition, the cross-correlation can be used to quantify time delays between two signals. We found that the maximum correlation occurred at a time lag of 0 minutes, indicating that changes in PhoP are quickly propagated to the downstream genes. While changes in GFP should, in principle, slightly lag changes in mCherry, measurement constraints, such as the temporal resolution of data collection, prevent the precise determination of the time of signal propagation. Additionally, the longer maturation time of mCherry in comparison to the GFPmut2 variant we used may reduce any apparent time lags.^48,59^ However, these constraints are universal across all experiments, meaning cross-correlations of the different gene reporters can be compared; we also note that these temporal correlations differ significantly from an unregulated constitutive control. These results indicate that there are no dramatic differences in response time between the tested PhoP regulon promoters, and that overall response times are short.

Although positive, lag-free cross-correlations imply that changes in input immediately induce an output response in the same direction, they do not capture the error of information transmission from input to output. In a low-error pathway, time-varying output signals can be readily attributed to distinct dynamic inputs. On the other hand, noisy signal transduction in the pathway would lead to overlapping outputs with high uncertainty in determining their input. To quantify the dynamic signal transduction reliability, we calculated mutual information between dynamic mCherry (input) and GFP (output) time series. Time-varying signals can encode more information than static readouts because they capture signal dynamics.^56^ However, mutual information decreases when time series data display high temporal overlap. We found that mutual information between mCherry and GFP time series for *phoP*, *pmrD*, and *slyB* was around 1.2 bits while *yrbL* had noisier signal transmission behavior with mutual information of 0.6 bits (Fig. 2D). Expectedly, with high overlap between its output time series, *yrbL* carried only half of the information exhibited by the other genes with clear, distinct outputs. Interestingly, compared to end-point, static flow cytometry measurements, *phoP* and *pmrD* exhibited higher input-output mutual information in dynamic analysis, indicating that coordinated temporal dynamics between input and output compensates for the overlap observed in static population measurements. Collectively, these results demonstrate that analyzing time-varying input-output relationships enables both characterization of temporal correlations between input and output as well as quantification of dynamic signal transmission fidelity.

### Joint dynamics of PhoP and downstream genes enable reliable encoding of input information

The differential signal transmission properties of downstream genes reflects the fact that regulatory pathways respond unequally to the same inputs. To examine how reliably regulatory molecules report information about incoming inputs, we set out to assess how accurately this information can be decoded. We assumed that optogenetic control places cells in one of three distinct input conditions, driven by the objective groups. High-fidelity signal transmission should enable accurate decoding that matches the imposed optogenetic condition, or the objective group, while noisy transmission would result in misclassification. Importantly, since upstream and downstream regulatory molecules both participate in signal transmission, we hypothesized that regulatory molecules jointly encode more reliable information than each alone.

To investigate the role of different downstream genes as indicators of input condition in response to external light inputs, we segregated the mCherry and GFP time series during the period of active control––i.e., hour 4 to 12––into the three control objective groups. We then compared both PhoP and downstream gene expression over time between the three objective groups (Fig. 3A). Here, rather than treating these as separate input-output signals, we examined their three-dimensional trajectories through time as integrated time series that collectively encode input condition information. Since PhoP and the downstream gene *jointly* report the input condition, we refer to these temporal trajectories ‘joint time series.’

**Figure 3.**
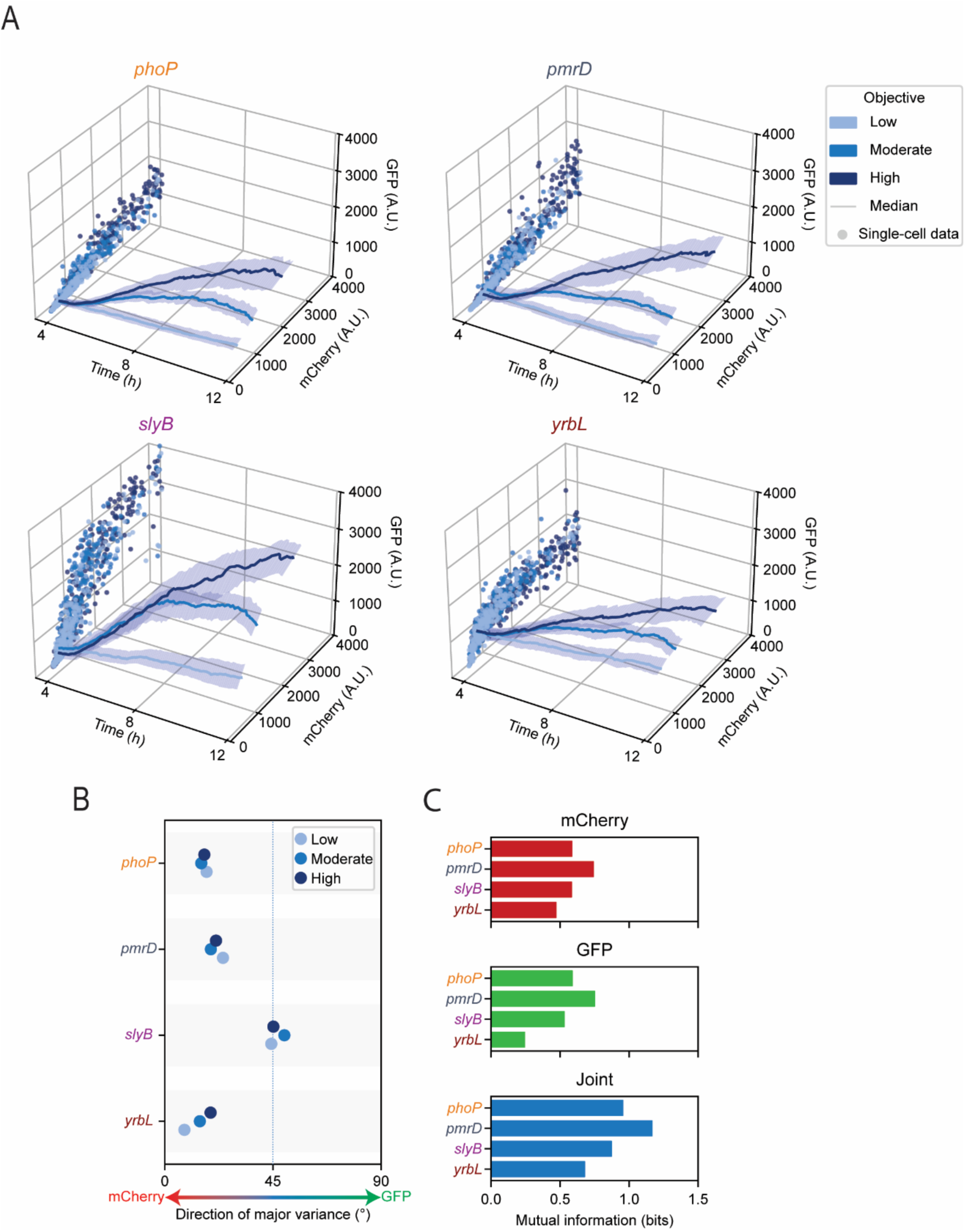
Heterogeneity in response to PhoP activation. (A) Median joint time series with mCherry and GFP. Scatter plot of projection of the joint time series onto mCherry-GFP plane is also shown. *n* > 200 cells for each objective. Error bands show 2^nd^ and 3^rd^ quintiles for each objective group over time. (B) The direction of major variance of the joint time series projection on the mCherry-GFP plane for each objective group per downstream gene. Lower angles correspond to higher variation along the mCherry (PhoP) axis while higher angles represent larger variation along the GFP (downstream gene) direction. (C) Mutual information between single or joint time series and the objective group.

The projection of joint time series on the mCherry-GFP plane illustrated the entire range of values that PhoP and downstream gene cover for each gene (Fig. 3A). While the low objective group was concentrated at low mCherry and GFP values (light blue points in Fig. 3A), inclusion of moderate and high objective groups allowed coverage to span a wider range of the mCherry-GFP plane. We found that each downstream gene exhibited a distinct joint time series for the three objective groups. For example, projection of the *slyB* joint time series demonstrates wide coverage of the mCherry-GFP plane, indicative of large variation of PhoP and *slyB* within each input condition. The differences in the projection patterns also reflect preferential changes in one dimension more than the other, revealing gene-specific patterns of coordinated variation. For example, the joint time series of *yrbL* mostly varies in the mCherry (PhoP) direction, indicating that *yrbL* has limited downstream sensitivity to changing inputs compared to *phoP*, *pmrD*, and *slyB*. To quantify the preferential variation of the joint time series, we calculated the first principal component of the joint time series projection on the mCherry-GFP plane (Fig. 3B). For each gene, the direction of the first principal component was largely consistent across the three objective groups, indicating that the gene-specific input-output relationship is input-invariant and persists across different input conditions. We found that *slyB* has balanced downstream sensitivity, displaying equal variation in both PhoP and *slyB* directions. On the other hand, *phoP* and *pmrD* had moderate downstream sensitivity with modest variation along the mCherry axis while *yrbL* showed low downstream sensitivity with dominant variation along the mCherry axis. Overall, these results underscore the variability of different joint PhoP-downstream gene time series in response to the same external inputs.

We also confirmed that the mCherry-GFP projection patterns were similar for each gene when measurements at a single time point were considered (Fig. S9). In addition, while individual mCherry-GFP projection patterns were gene-specific, temporal as well as cell-to-cell heterogeneity in joint time series among the cells of a single population could be observed. To illustrate this heterogeneity, we plotted the ratio of GFP to mCherry over time for different downstream genes for 25 randomly selected cells (Fig. S10). This simplified illustration of mCherry-GFP projections over time elucidates the heterogeneity in joint time series within the same population.

In addition to different sensitivity, genes exhibited varying temporal discriminability. For instance, the joint time series of *pmrD* clearly separate from each other early during the 4-12 hour period while the joint time series of moderate and high objectives of *slyB* have high overlap with each other. Overlapping joint time series indicates poor discrimination of input conditions by the regulatory pathway. To investigate the reliability of integrated PhoP-downstream gene temporal signals as determinants of input condition, we calculated the mutual information between the joint time series and the input group for each gene. This quantity implies the certainty of determining the input condition, given the history of PhoP and downstream gene expression through time. As opposed to mutual information analysis of static signals (Fig. 1G and H) where a snapshot of PhoP and downstream gene measurements were used to attribute one of the four input groups to each fluorescence population, here, the gene expression history in single cells is assumed to be a multivariate signal encoding the information of the input condition.

Since each gene displayed a distinct pattern of input-output covariation, we hypothesized that the joint dynamics of PhoP and each downstream gene would carry more information about input condition than either signal alone. We observed that individually, mCherry and GFP time series resulted in low mutual information values below 1 bit (Fig. 3C), reflecting poor discrimination. Specifically, mutual information extracted from GFP time series was significantly lower for *yrbL* given its low downstream sensitivity, indicating that determining the objective group based on the *yrbL* signal is unreliable. However, the mutual information increased substantially when the joint time series were considered as reporters of the input condition. For *pmrD*, this value rose above 1 bit. For reference, 1.58 bits corresponds to perfect discrimination between the three input groups (log_!_ 3). Imperfect discrimination can be attributed to temporal and cell-to-cell heterogeneity in expression of PhoP and downstream genes that inevitably introduces noise and uncertainty to the input condition discrimination. While this noise makes determining input conditions uncertain, it may be a bet-hedging mechanism that makes some cells tolerant to potential future challenges. Collectively, this analysis shows that the joint dynamic PhoP-downstream gene signals enable a multivariate information encoding mechanism that enhances the reliability of input condition detection compared to what either signal provides alone.

### Role of PhoP and PmrD in Polymyxin B tolerance

The history of gene expression inside a cell collectively determines cell state.^31^ This cell state reflects the cell’s current physiological condition and predicts its likelihood of growth or survival in the face of environmental changes. By monitoring temporal variation of multiple signaling molecules, cells perceive their state and adaptively respond to changes in the environment. However, different signaling molecules participate unequally to this growth- or survival-relevant state perception.^31^ Inspired by this multimodal cell state perception mechanism, we wanted to investigate whether joint time series of PhoP and downstream gene expression could predict cell survival.

To understand how gene-specific transmission properties contribute to cellular decision-making (*e.g.*, in the face of an external challenge), we asked how PhoP and downstream gene expression jointly encode cellular survival information through their coordinated temporal dynamics. To answer this question, we investigated the functional consequences of PhoP system activation by challenging cells with lethal concentrations of the antimicrobial peptide Polymyxin B.

Polymyxin B is a short, cationic lipopeptide used to treat infections of gram-negative bacteria. Its strong bactericidal effect and efficacy against multidrug resistant bacteria make it an important drug in clinical settings.^60^ Polymyxin B targets bacterial membranes as the primary mode of action, making outer membrane modifications the main strategy for resistance.^61^ A homolog of the PhoP family gene *pmrD* is strongly implicated in Polymyxin B resistance in *Salmonella enterica*, where it plays a role in linking PhoP activation to activation of PmrAB, a two-component regulatory system responsible for membrane modification.^25^ However, the role of *pmrD* in *E. coli* is less clearly defined. While *pmrD* was previously thought not to connect *phoPQ* and *pmrAB* expression in *E. coli,*^43^ a later study found evidence that *pmrD* can activate *pmrA* under certain conditions, such as during the low Mg^2+^ conditions which also lead to high activation of PhoPQ. It was additionally found that while *pmrD* is typically upregulated by PhoPQ, it may also be upregulated in a PhoPQ-independent manner.^24^ Thus, the precise relationship between PhoP, expression of *pmrD,* and the effect of each gene in enabling Polymyxin B resistance remains uncertain. This motivated our focus on PhoP and *pmrD* for our analysis on Polymyxin B survival. In particular, we sought to characterize the extent to which PhoP, *pmrD*, or their joint variations resulted in protection against Polymyxin B.

To investigate the role of *pmrD* in Polymyxin B resistance in *E. coli*, we observed cells containing the *pmrD* reporter in the mother machine before and after switching to media containing lethal concentrations of Polymyxin B (500 ng/mL, Fig. S11). We again initiated low, medium, and high levels of PhoP activation, following a four-hour equilibration period. Polymyxin B was added to the growth media starting at hour eight (Fig. 4A). In contrast to the previous set of experiments, the control strategy was maintained for the duration of the experiment. This resulted in populations with low, medium, and high levels of PhoP and *pmrD* expression (Figs. 4A and 4B). We note that control accuracy was impacted after the addition of Polymyxin B, when cells begin to grow more slowly, filament, or die.

**Figure 4.**
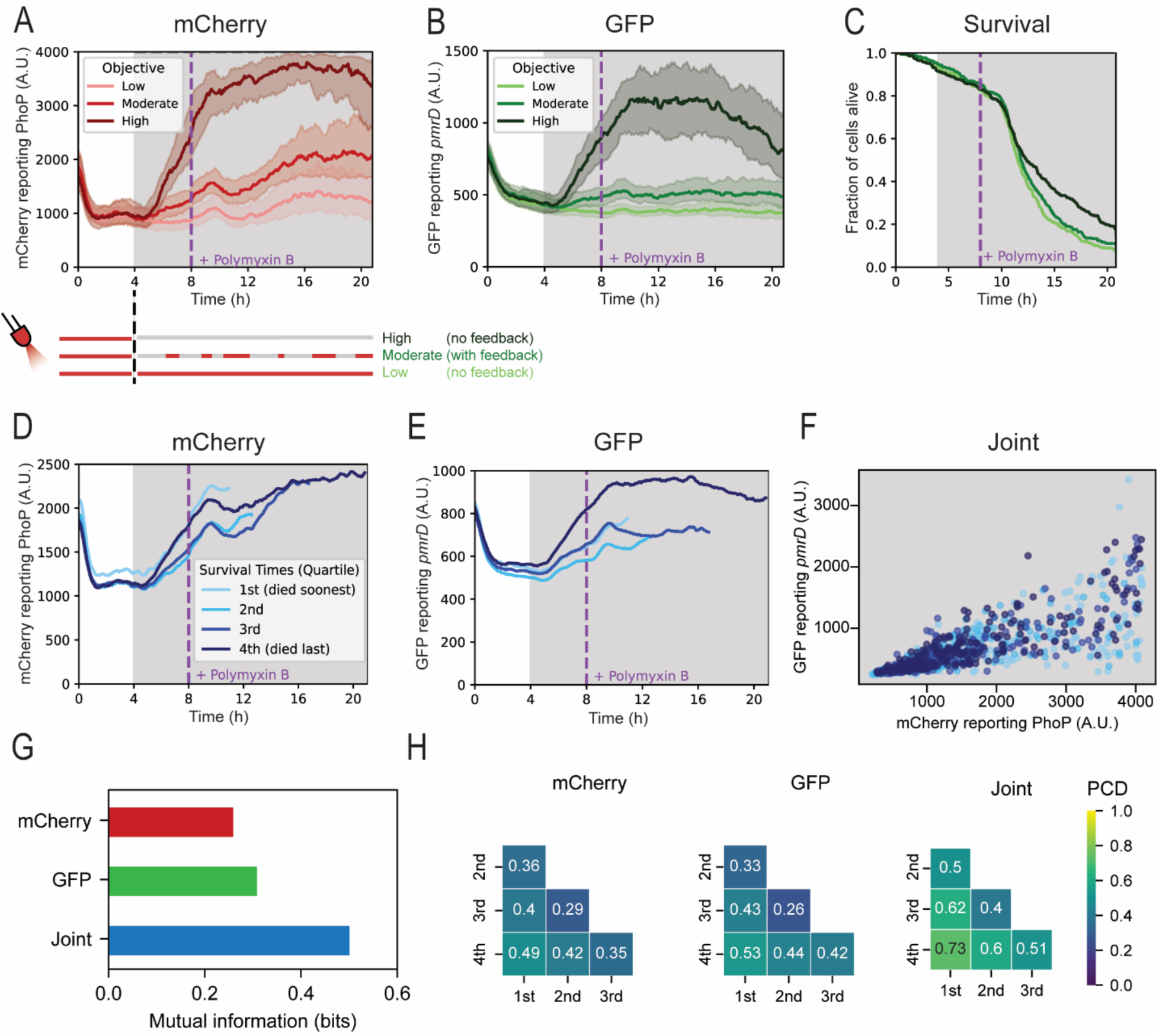
Role of *pmrD* in protecting cells from the antimicrobial peptide Polymyxin. **B.** (A) mCherry reporting PhoP levels for all cells over time in low, moderate, and high objective groups. The specified objective group begins at time t = 4 hours and continues until the end of the experiment. At time t = 8 hours, Polymyxin B is added to the cell media at a final concentration of 500 ng/mL. (B) GFP levels, from the *pmrD* reporter, for cells in each objective group over time. Median plus middle quintile shown for each objective group. *n* > 200 cells per objective. (C) Fraction of cells remaining alive for each objective group over time. (D) Mean mCherry over time for cells from all objective groups. Cells are partitioned into quartiles based on time of death after Polymyxin B exposure, with the 1^st^ quartile representing the shortest-surviving cells and the 4^th^ quartile representing the longest-surviving cells. Note that the durations of the data traces vary between the four groups due to differences in time of death. (E) Mean GFP from the *pmrD* reporter over time for all cells. (F) Scatter plot of joint time series projection on mCherry-GFP plane from hour 4 to 8 of the experiment. (G) Mutual information between the single or joint time series and the survival quartile. Dynamic signals from hours 4 to 8 of the experiment were used for computation of mutual information. (H) Pairwise probability of correct discrimination (PCD) between survival quartiles extracted from single or joint time series.

Throughout the experiment, we tracked cell growth rate and used this information to determine when growth ceased, at which point cells were classified as dead (Fig. 4C). We found that a subset of cells die in the hours prior to Polymyxin B exposure due to cell aging or spontaneous filamentation, which can occur naturally as cells accumulate damage over time while growing in the microfluidic device.^57^ After Polymyxin B exposure, cells die more quickly, with fewer than 20% of cells remaining alive by the end of the experiment.

To probe whether either PhoP, *pmrD*, or the joint time series encode enough information to predict cell survival, we divided the cells into four survival groups. Unlike our previous analysis, where we segregated the mCherry (PhoP) and GFP (*pmrD*) expression based on the three objective groups, here, we divided the time series data based on cell survival quartile. Dividing the survival data into quartiles provides four biologically interpretable groups ranging from ‘most susceptible’ to ‘most resistant’ cells while maintaining a reasonable sample size per group for robust information theoretic analysis. Each quartile has a distinct cellular fate, with the 1^st^ survival quartile including the cells that survived the shortest amount of time after Polymyxin B exposure and the 4^th^ including the cells that survived the longest. We hypothesized that the history of PhoP and *pmrD* expression until the moment of addition of Polymyxin B would be strong predictors of cell survival.

Population analysis of survival quartiles indicated balanced representation of cells from each input objective per survival quartile, although the cells controlled with the high expression objective were slightly overrepresented among the longest-surviving cells (Fig. S12). All four survival groups experienced a similar rise in PhoP (mCherry) signal upon initiation of active control at hour 4 (Fig. 4D). This further confirms that survival groups do not necessarily correlate with the objective groups (Fig. S12). Similarly, *pmrD* (GFP) expression generally increased for all survival quartiles after optogenetic control began, with the 4^th^ quartile displaying the largest increase (Fig. 4E), suggesting that *pmrD* time series are likely a stronger predictor of cell survival than PhoP dynamics. However, investigation of joint PhoP-*pmrD* space revealed variation along both PhoP and *pmrD* axes for all survival groups (Fig. 4F), highlighting the heterogeneity of PhoP and *pmrD* expression within the survival quartiles. This temporal expression heterogeneity consequently leads to representation of cells from all optogenetic objective groups among the longest-surviving cells. These results suggest that high *pmrD* expression is a strong factor in survival with a potential role for PhoP in enhancing the likelihood of predicting survival. Simultaneously, the presence of cells from low and moderate optogenetic objectives among the longest-surviving cells with high *pmrD* expression indicates the role of heterogeneous gene expression in increasing survival likelihood.

To investigate the role of joint time series in predicting cellular survival, we calculated the mutual information between the survival group and dynamic PhoP, *pmrD*, or joint time series data. In contrast to prediction of input condition from dynamic signals, where the PhoP signal universally exhibited stronger performance than downstream genes, here, mutual information of *pmrD* and survival was slightly higher than PhoP (Fig. 4G). This is aligned with the observation of distinct *pmrD* dynamic signals for the 4^th^ survival quartile and further verifies the role of *pmrD* in conferring tolerance to Polymyxin B. However, a notable increase in mutual information is achieved by the joint PhoP-*pmrD* time series, indicating that knowing the history of expression of both PhoP and *pmrD* prior to exposure to Polymyxin B leads to stronger predictions of cell survival. This enhanced predictive power is not due to a larger number of variables for joint time series data compared to single PhoP or *pmrD* signals. We confirmed this by showing that concatenation of identical PhoP or *pmrD* as well as random shuffling of joint time series led to reduced mutual information (Fig. S13), thus joint PhoP-*pmrD* time series carry meaningful information. Importantly, this means that the interplay of PhoP and *pmrD*, rather than PhoP or *pmrD* solely, determines cellular fate. Considering the stronger predictive power of joint time series in determining input conditions, these results underscore the multimodal mechanism by which signaling molecules determine cell state and fate.

In addition to mutual information, our analysis offers quantitative insight into pairwise discrimination between survival quartiles. This metric, known as probability of correct discrimination (PCD), quantifies the likelihood of correctly discriminating between two survival quartiles, ranging from 0% (random chance) to 100% (perfect discrimination). Since PhoP dynamic signals exhibited large overlap between the quartiles, the PCD was low for all comparison groups with the lowest being only 35% likelihood of correctly discriminating between 3^rd^ and 4^th^ quartiles. While *pmrD* signals enabled modestly higher likelihoods of correct discrimination, the joint time series outperformed single time series for all comparison groups with the highest PCD of 73% between 1^st^ and 4^th^ quartiles. Collectively, these results highlight the stronger predictive power of joint time series as multivariate proxies of cell state in both determining cell fate as well as correctly predicting the duration of cell survival. This adds additional evidence that while PhoP and *pmrD* both have a protective effect on cells encountering antimicrobial stress, neither alone is solely predictive of cellular survival, and it is their interplay that may matter most for conferring tolerance.

## Discussion

While heterogeneity in the expression of stress response transcription factors has been characterized in a wide range of contexts, few studies have examined how this heterogeneity is propagated to the downstream genes that enact stress tolerance. We tested a select set of PhoP-regulon genes and observed an increase in their expression as PhoP levels increased. However, the response patterns were widely variable and distinct for each gene. Different target genes required different amounts of PhoP for activation and had different fold-changes when maximally induced.

Further, the fidelity of signal propagation from PhoP to downstream genes was found to vary widely, with some genes displaying high-quality signal transmission while others exhibited noisy signal transduction. We also found that, analogous to stress response genes which exhibit seemingly stochastic patterns of high and low activity over time and across populations of isogenic cells, individual cells may also exhibit heterogeneous downstream responses to an identical upstream input. These results highlight the advantages of examining multiple genes in a pathway, especially when seeking to link heterogeneity in gene expression to functional outcomes such as survival.

Our results underscore related work showing the differential effect of identical PhoP levels upon a range of gene promoters in its regulon.^23,27,62^ While these differences in gene network signal propagation partly stem from differences in overall PhoP levels and the tendency of some promoters to saturate, we also noted cell-to-cell heterogeneity that points to the role of noise or other intracellular factors in determining the PhoP response.

Importantly, our live, single-cell microscopy platform allowed us to investigate dynamic signal transmission properties of the PhoP regulon. Unlike previous studies,^53–55^ here, we simultaneously captured the temporal heterogeneity in expression of both PhoP and its downstream regulated genes in response to dynamic optogenetic inputs. This allowed us to uncover the transmission reliability of dynamic signals for downstream genes, providing further insight into differences in responsiveness and sensitivity across genes within the PhoP regulon.

Additionally, by monitoring both upstream and downstream signaling molecules, we demonstrated the enhanced predictive power of joint, dynamic signals in determining external inputs as well as cellular fate. This finding aligns well with the proposed model of multimodal cell state perception demonstrated for human epithelial cells responding to epidermal growth factor signals.^31^ While our microscopy platform allowed for simultaneous measurement of two signaling molecules, we note that monitoring expression levels of more signaling molecules involved in the PhoP regulon or another stress-response pathway would shed further light on the ability of expression history of bacterial signaling molecules to predict cell state and survival.

We have shown that even subtle heterogeneity in PhoP regulon gene expression can have important consequences for cell survival. Our experiments indicate that in conditions of non-saturating PhoP activation, some cells can gain an edge in Polymyxin B survival due to the combined effects of their PhoP expression history and disproportionately high expression of *pmrD*. While the increase in *pmrD* may not be solely caused by PhoP, our findings suggest that the interplay of both PhoP and *pmrD* leads to higher chances of cell survival. This suggests that the upstream regulator and functionally relevant downstream signaling molecules act together to prepare the cell for potential future challenges. Part of the reason for these results may also be that *pmrD* can be activated in a PhoP-independent manner under certain conditions, including low Mg^2+^.^24^ These results point to potential gene targets that could be disrupted in combination with Polymyxin B treatment, or tested in clinical isolates that show resistance to the antimicrobial peptide for further insight into resistance mechanisms.^63^ Despite PhoP being the main regulator of a range of virulence traits, precision targeting of *pmrD,* alone or in tandem with PhoP, may prove more effective in characterizing or preventing Polymyxin B tolerance.

Possible extensions of this work include testing other functional outcomes of variations in PhoP activity, such as the ability of cells to withstand acid or osmotic stress or to tolerate exposure to other antimicrobial peptides, since PhoP is involved in coordinating a wide array of stress tolerance mechanisms. While we chose to focus on exploring the relationship between total PhoP activity on downstream promoters, alternate experimental setups could be used to further disentangle the role that native genetic feedback mechanisms play in the system, where both positive and negative feedback loops can exhibit variable behavior depending on the strength of PhoP regulon activation.^64–66^ This could be accomplished by leaving the native *phoP* gene intact, incorporating methods for tracking PhoP phosphorylation dynamics in real-time,^67^ or employing sophisticated feedback compensation techniques made possible by optogenetic control.^68^

While the downstream genes selected in this study are directly regulated by PhoP, examining more complex pathways consisting of signaling cascades and various feedback motifs will elucidate information processing capabilities and the role of noise and heterogeneity in complex signaling networks. From a technical perspective, utilizing fast-maturing fluorescent proteins would allow for capturing the nuances of dynamic signal transmission, thus allowing more accurate measurements of any transmission delay. Measuring short-lived dynamics would additionally enable capturing dynamic downstream responses to time-varying inputs, such as oscillating PhoP levels.

Protein-level accumulation of gene products in the PhoPQ regulon could also be examined, as this has been shown to lead to additional heterogeneity in the response of other systems.^69^ Similar studies could also be carried out with other transcription factors implicated in cell stress to uncover broader patterns in gene regulation. Together, these studies will shed light on the complex and sometimes subtle ways in which heterogeneity in gene expression associated with stress response can impact cell survival or evasion of antimicrobial therapies.

## Materials and Methods

### Plasmid and strain development

All experiments were carried out using the *E. coli* BW25113 strain or its derivatives. The BW25113 Δ*phoP* knockout strain was sourced from the Keio collection.^70^ For this strain, we excised the kanamycin resistance cassette using the pCP20 plasmid,^71^ and the pCP20 plasmid was subsequently cured by growing cultures at 43°C overnight. To confirm curing of both kanamycin resistance and the pCP20 plasmid, we tested single colonies to verify sensitivity to both kanamycin and ampicillin. We constructed the variant of the *phoP* gene that was used in all experiments (unless otherwise noted) by introducing the point mutations described in Miyashiro *et al.*^27^ This variant behaves as though constitutively phosphorylated. The BW25113 Δ*phoP*:P*_cpcG2_*-*phoP* and BW25113:P*_cpcG2_*-*tetR* strains were constructed using lambda red combination.^71^ All cassettes were integrated into the locus downstream of the *nupG* gene, as previously described,^14,72^ using the homology regions 5’-TCGTGTCCCGACAGGCACACAGACGGTTAGCCACTAATTA-3’ and 5’-AAAAAAACGGGTCACCTTCTGGCGACCCGTTTTTCTTTGCG-3’.

All *GFPmut2* reporter plasmids were sourced from the collection developed by Zaslaver *et al.*^48^ All promoter sequences were verified by Sanger sequencing (Quintara Bio). We noted that the promoter sequence of the *nagA* reporter in the Zaslaver collection did not contain the *nagA* promoter sequence reported in RegulonDB.^73^ Therefore, we cloned the *nagA* promoter sequence indicated in RegulonDB upstream of *GFPmut2* using Gibson assembly through two sequential PCR reactions with forward primers ATATTGCATTTAACGAACCGGCGTCTTCTCTCGAGGGGATCCTCT and GCGGTGTAGGTAACGACGGTCATATTGCATTTAACGAACCG, to add the *nagA* promoter as the 5’ overhang to *GFPmut2*, as well as the reverse primer for both: CCTTTCGTCTTCACCGTTCTTACGGAAAAATTCATCTGTTTATGG. The reporter plasmid used in TetR control experiments (described below, Fig. S14) consisted of pBbS2k with TetR excised and with the gene encoding mCherry replaced with *sfGFP*.^74^ The constitutive reporter was constructed by replacing the promoter sequence with a modified version of the phage T7 A1 promoter (5’ - TTATCAAAAAGAGTATTGTCTTAAAGTCTAACCTATAGGAAAATTACAGCCATCGAGAGGGACACGGCGAA - 3’).

mCherry was expressed under the control of the P*_cpcG2_* promoter from a separate plasmid, which consisted of the pSR58.6 plasmid with *mCherry* replacing *sfGFP*. This plasmid was transformed along with pNO286-3 and the relevant reporter plasmids into each strain. This resulted in a system with a high dynamic range of mCherry expression, which allowed it to serve as an easily measurable readout of system activation while TetR or PhoP were modulated at lower levels. pSR58.6 and pNO286-3 correspond to Addgene plasmids #63176^75^ and #107746^76^, respectively, from Dr. Jeffrey Tabor’s lab.

### Bulk culture optogenetic experiments

All bulk culture optogenetic experiments were carried out using an optoWELL-24 optogenetic well plate device (Opto Biolabs GmbH). Overnight cultures were diluted 1:100 into fresh lysogeny broth (LB) (Fisher) and 600 μL total volume was added to each well of a 24 well plate. The optoWELL-24 was programmed with individual red (657 nm peak intensity) and green (519 nm peak intensity) light intensities for each well. Each well received red light at an intensity of 0.36 mW/cm^2^, while green light varied in intensity from 0 – 0.208 mW/cm^2^ as detailed in the figure legends. The well plate was placed inside the optoWELL-24 device, and the device was placed in a shaking incubator at 37°C and 250 rpm. After 5-6 hours, a BioTek Synergy H1m plate reader was used to measure optical density (A_700_), GFP (485 nm excitation, 528 nm emission), and mCherry (580 nm excitation, 610 nm emission) for each well. Unless otherwise noted, optical density was measured at 700 nm to reduce overlap with the mCherry emission spectrum. 15 μL was also sampled from each well and diluted into 135 μL of phosphate-buffered saline for flow cytometry using a GuavaFlow easyCyte HT flow cytometer. 5000 flow events were acquired for each sample. Flow cytometry data were gated to remove dead cells and cells with non-positive fluorescence values using the FlowCal^77^ package in Python. Fluorescence distributions were fit to a kernel density estimation curve using the Seaborn package in Python, with a grid size of 50.

Geometric means of fluorescence measurements for each reporter and stimulation level were used to fit a Hill equation:

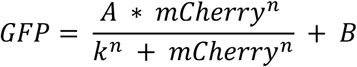

The parameters 𝐴, 𝑘, 𝑛, and 𝐵 were independently fit for each reporter strain using the curve_fit() function from the SciPy package in Python.^78^ Fit values are listed in Table S1.

To validate that our optogenetic platform was not introducing spurious effects on expression from gene reporters, we performed additional bulk culture experiments using a synthetic gene inverter circuit. A *gfp* reporter was placed under control of the P_Tet_ TetR-repressible promoter, while the optogenetic CcaS/R system was used to control TetR repressor levels by placing the *tetR* gene and the *mCherry* gene downstream of the P*_cpcG2_* promoter (Fig. S14A). We verified that increasing levels of green light stimulation resulted in increasing levels of mCherry, while GFP levels decreased (Fig. S14B). Overall, the system showed a consistent inverse relationship (Fig. S14C), adding confidence that our platform could detect input-output relationships.

### Quantification of gene expression with qPCR

RNA purification, RT-PCR, and qPCR experiments were performed following conventional protocols.^79^ Briefly, overnight cultures grown in the dark were diluted 1:100 into 1 mL LB and were dispensed in a 24 well plate. The cultures were grown while exposed to the designated green light intensities as described in the previous section. After a 6 hour incubation, the optical density (A_600_) of all cultures was measured, and the culture optical density was normalized to ∼0.5. Upon normalization, a 600 µL sample from each culture was mixed with 600 μL of a chilled 1:1 mixture of ethanol:acetone and samples were stored at -80 °C overnight. Cells were thawed on ice, centrifuged, harvested, and mixed with 100 μL 1 mg/mL Lysozyme from chicken egg (Sigma Aldrich). Total RNA was next extracted from cell lysates using a Monarch Spin RNA Isolation kit (New England Biolabs). cDNA synthesis was then performed on whole RNA extract using random hexamer primers with the GoScript Reverse Transcriptase kit (Promega). Lastly, the abundance of target cDNAs, namely *phoP*, *mCherry*, *slyB*, *gfp*, and the housekeeping gene *gapA*, were quantified through a KAPA SYBR FAST (Roche) qPCR kit using a Bio-Rad CFX96 real-time PCR detection system. qPCR primers used for amplification of each target cDNA are listed in Table S4. To quantify the normalized expression level for each target gene, the ΔC_q_ method was followed. The C_q_ level for each target gene was calculated using the Bio-Rad CFX Maestro software, and the normalized expression level was then calculated using:

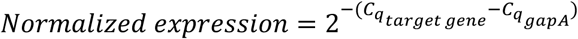

### Mother machine microfluidic device

Mother machine devices were produced as previously described.^14^ Briefly, SU-8 resin molds were made via stereolithography based on the design available at https://gitlab.com/dunloplab/mother_machine. Polydimethylsiloxane (PDMS) (Dow Corning Sylgard 184) was poured into the molds, cured overnight at 75°C, and cut from the molds. Inlets and outlet holes were punched at either end of each channel using a 0.75 mm biopsy punch and the chip was plasma bonded to a glass slide. Following binding, chips were returned to the 75°C incubator for an additional 24 hours before use.

### Time-lapse microscopy

Overnight cell cultures were refreshed 1:100 into fresh LB medium and grown for 2-3 hours before being loaded into the mother machine microfluidic device. The device was spun at 5000 rpm for 3 minutes to load cells and then media tubing was connected at the inlet and outlet ports. LB supplemented with 2 g/L Pluronic F-127 (to reduce cell adhesion to the device) was supplied to the chip at a rate of 20 μL/min using a peristaltic pump. For all mother machine experiments, a concentration of 50 μg/mL chloramphenicol was provided as the sole antibiotic for plasmid maintenance. This concentration was selected to minimize cell stress while still preventing media contamination.

Cells were imaged with the following order and exposure times: phase contrast (50 ms), mCherry (35 ms), and GFP (50 ms). Following imaging, cells were either exposed to 100 ms of red light from a digital micromirror device (DMD) in order to repress CcaS/R activity, or they received no DMD stimulation. Additionally, the full chip was exposed to low levels of constant red light from an Arduino-controlled NeoPixel Ring (Adafruit 1586) set to 25% maximum intensity, which was affixed directly above the microscope platform, in order to reduce the strength of background activation.

### Deep model predictive control

For the moderate light exposure case, deep model predictive control was enacted on cells using the platform previously developed.^58^ Unless otherwise specified, control parameters used were the same as in the original study. Briefly, snapshots of the cells were segmented and features such as fluorescence and cell length were extracted in real-time.^80^ The features were fed into the deep model predictive control algorithm, which uses a trained model to predict which optogenetic stimulations are most likely to make the cell’s mCherry fluorescence match the desired levels over time. The optimal optogenetic stimulations were used to create an image that was then loaded into the DMD to stimulate individual cells. We acquired a new training set for the prediction model by subjecting the optogenetic strains to random red stimulations using the DMD. Three experiments were conducted to acquire this training set, each tracking approximately ∼2,000 cells over a period of at least 16 hours. Each training set contained cells with several different PhoP-responsive GFP reporter plasmids, although in principle the specific reporter used should not matter, because the model was not trained on GFP values and only uses mCherry fluorescence values.

### Mother machine experiment data analysis

Cell segmentation and feature extraction were carried out on-the-fly during experiments using DeLTA software,^80^ as previously described.^58^ This provided data on fluorescence and cell area for each mother cell over the course of the experiment. Growth rate was calculated *a posteriori* from cell area data, according to the following formula:

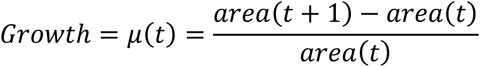

To simplify analysis, cells experiencing division or non-physiological growth rates in any given frame (namely, rates < -0.2 hr^-1^ or > 0.3 hr^-1^), which occasionally occurred due to segmentation algorithm glitches or cell death, were assigned “NaN” values. “NaN” values were also assigned to any objects identified by the segmentation algorithm whose total area was too small to represent a true cell. Growth rate data was then smoothed with a moving average filter over 5 frames to remove artefacts.

After growth rate was calculated over time for all cells, dead cells were filtered out from the dataset in order to draw conclusions based only on living cells, unless otherwise specified. Cells were removed if they had been assigned a “NaN” growth rate or a growth rate below 0.1 hr^-1^ for four consecutive frames. This metric was chosen to avoid removing cells which had only momentary slow growth or which experienced segmentation glitches for a small number of frames. Only cells which survived for the full length of the experiment were considered for analysis. For each experiment, at least 100 surviving cells were present for each reporter and objective group.

Normalized cross-correlations were calculated by using the correlate function from NumPy, then dividing by the square root of the product of each signal’s auto-correlation with itself.

### Polymyxin B minimum inhibitory concentration (MIC) assay

Overnight cultures of wild-type BW25113 cells were diluted 1:100 in M9 minimal media containing M9 salts, 2 mM MgSO_4_, 0.1 mM CaCl_2_, and 0.4% glucose. Cells were grown for 4 hours at 37 °C in a shaking incubator. Meanwhile 120 µL minimal M9 solution supplemented with a serial dilution of Polymyxin B sulfate (MedChemExpress) was dispensed in a 96 well plate. After incubation, 5 µL of cell culture were added to each well, and the well plate was incubated for 18 hours at 37 °C in a shaking incubator. After the incubation, the optical density (A_600_) was measured using a BioTek Synergy H1m plate reader.

### Polymyxin B survival experiments

Polymyxin B survival experiments were carried out using a similar protocol for optogenetic stimulation, with the following differences: After the first four hours of red light DMD stimulation, the step-on control objectives were maintained for the full course of the experiment, rather than stepping-off at hour 12. Additionally, after hour 8, Polymyxin B sulfate was added to the cell media at a final concentration of 500 ng/mL. Instead of removing cells which did not survive for the full duration of the experiment from analysis, all cells which survived at least until the addition of Polymyxin B to the media were included in the analysis but were then sorted into quartiles based on the time of cell death. A cell was marked as dead if the cell had a growth rate below 0.1 hr^-1^ for four consecutive frames, as described above. The frame in which this occurred was noted for each cell. Cells were then sorted into quartiles based on this frame of death, with the 1^st^ quartile representing cells that survived the shortest, and the 4^th^ quartile representing cells that survived the longest.

### Statistical analysis

Pairwise Jensen-Shannon divergence (JSD) was calculated for the mCherry and GFP distributions presented in Fig. 1C using custom Python code. JSD is the average of two Kullback-Leibler (KL) divergences between each distribution and the mean distribution of the pair:

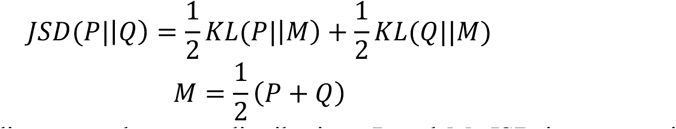

Where KL(P||M) is the KL divergence between distributions P and M. JSD is symmetric and bounded between 0 (for completely identical distributions) and 1 (for fully distinguishable distributions). Since we were interested in comparing the shape and spread of distributions and not their distance from each other, we centered all populations around zero by subtracting the mean from the data. The pairwise JSD was then calculated through kernel density estimation of each population followed by calculation of KL divergence.

Coefficients of variation (CVs) for GFP populations were computed using the standard deviation and mean functions from NumPy. To calculate statistical significance between CVs across objective groups, 1000 bootstrap resampling iterations with replacement were performed. For each iteration, the original data within each group were resampled, the CV for each group was computed, and the difference in CVs was calculated. The *p*-value was then estimated as the proportion of bootstrap iterations in which the resampled CV difference exceeded the observed CV difference under the null hypothesis of no difference. To compare correlation coefficients, individual correlation coefficients were Fisher transformed to Z values, and their difference was tested for statistical significance.

### Information theoretic analysis

Here, we summarize methods of calculating mutual information. Further details are presented in Supplementary Information. All custom analysis codes are available on GitLab at https://gitlab.com/dunloplab/phop_regulon-information_theory.

Maximum mutual information or channel capacity between discrete input groups and mCherry or GFP response populations were calculated following the procedure described in Hansen and O’Shea^53^ and Lee *et al.*^54^. Channel capacity is defined by the following equation:

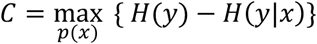

Where *H(x)* is Shannon information entropy. Through the definition of information entropy, channel capacity can be written as:

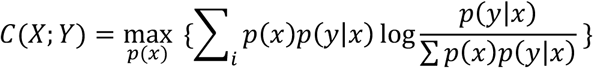

Where *p(y|x)* stands for the probability distribution of the mCherry or GFP population, obtained through binning, and *p(x)* is the optimal marginal distribution of inputs. Custom Python code was written to find the optimal input distribution using the iterative Blahut-Arimoto algorithm.^81,82^ To remove the bias due to bin size, discretization was performed for bin numbers ranging from 5 to 50. The unbiased channel capacity was calculated as the average of channel capacities for bins between 20 and 50.

To compute the mutual information between continuous uni- and multivariate signals presented in Fig. 1H and Fig. 2D, respectively, a non-parametric method named Kraskov-Stögbauer-Grassberger (KSG)^83^ was utilized. KSG relies on k-nearest neighbors to estimate joint and marginal distributions of input and output and does not suffer from bias due to the bin size. Mutual information was calculated using:

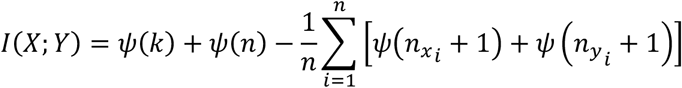

Where k, n, 𝑛_*x_i_*_, and 𝑛_*y_i_*_ are the number of nearest neighbors, sample size, and number of neighbors in marginal x and y spaces. Additionally, 𝜓(𝑥) is the digamma function. For Fig. 2D, only the segment corresponding to active control during the experiment (*i.e.,* hours 4 to 12) of the dynamic multivariate mCherry and GFP signals was used to calculate mutual information.

Lastly, mutual information between discrete input (or objective) groups and dynamic multivariate signals presented in Fig. 3C as well as 4G was calculated using the Statistical Learning-based Estimation of Mutual Information (SLEMI) R package.^84^ SLEMI fits a logistic regression model to the data for classifying multivariate data into distinct classes. The logistic regression model generates classification probabilities *p(x|y)* for each dynamic signal. These probabilities can then be used to calculate mutual information through the following equation:

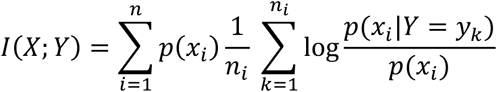

Where 𝑛_*n*_ is the sample size for each objective group. Data were segregated based on the input group and a custom R code was used to calculate mutual information with the SLEMI mi_logreg_main function. In addition, the probability of correct discrimination (PCD) was calculated using the built-in prob_discr_pairwise function in SLEMI. For each dynamic signal *y*, PCD is defined as the following:

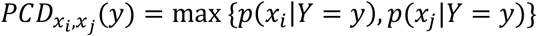

Where *p(x_i_|y)* is the probability of classifying signal *y* under the *i*th group. Pairwise PCD between input groups is then calculated by averaging the PCDs for all signals belonging to the two input groups:

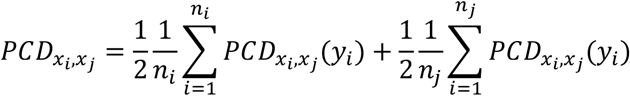

Since random choice between two groups has 50% probability, the PCD was transformed in order to have an intuitive range from 0 to 100% corresponding to totally random or completely certain discrimination, respectively.

## Acknowledgements

We thank Dr. Razan Alnahhas and Dr. Virgile Andreani for helpful discussions. Dr. John Ngo and Dr. Heidi Klumpe provided useful input on the manuscript. This work was supported by NSF grants MCB-2324909 and MCB-2032357 and NIH grant R01AI102922 to MJD. CMB received support from the NSF Graduate Research Fellowship under grant DGE-1840990.

## Author contributions

C.M.B and M.J.D. conceived and designed the study. J.-B.L. developed the optogenetic hardware and software platforms. H.M. conducted the information theory analysis. C.M.B., H.M., and F.J. performed experiments and conducted data analysis. C.M.B., H.M., and M.J.D. wrote the manuscript with input from J.-B.L. and F.J.

