## Supplementary Information for "Dynamic heterogeneity in an *E. coli* stress response regulon mediates gene activation and antimicrobial peptide tolerance"


1. Supplementary Methods
2. Supplementary Figures
3. Supplementary Tables

**<u>Supplementary Methods</u>**

***Information theoretic analysis***

Mutual information was calculated following Shannon's definition (Shannon, 1948):

$$I(X;Y) = H(Y) - H(Y|X)$$

in which *X* and *Y* stand for input and output signals and H(X) represents the information entropy of variable *X*. By expanding the term representing information entropy, the mutual information can be written as:

$$I(X;Y) = \int p(x,y)\log\frac{p(x,y)}{p(x)p(y)}\,dy\,dx$$

where *p(x,y)* is the joint probability density function of *X* and *Y* while *p(x)* and *p(y)* represent marginal distributions of *X* and *Y*, respectively.

***Mutual information of static readouts***

In Fig. 1C, the green LED intensity can be considered as the input while mCherry or GFP values are taken as the output. In this case, the measured fluorescence values represent the samples from the conditional probability *p(Y|X)* distribution where *X* is the input (LED power intensity). Since the input is discrete here, calculating maximum mutual information (MI, also known as channel capacity) is more tractable compared to mutual information since the marginal distributions of *X* and *Y* are unknown. To calculate MI, an iterative optimization algorithm known as Blahut-Arimoto (Arimoto, 1972; Blahut, 1972) which finds an input distribution that maximizes the mutual information was used. Therefore, MI here is defined as:

$$MI = \max_{p(x)}\big(I(X;Y)\big) = \max_{p(x)}\Big(\int p(x)p(y|x)\log\frac{p(y|x)}{\int p(y|x)p(x)\,dx}\,dy\,dx\Big)$$

To estimate the conditional probability distribution *p(y|x)*, the flow cytometry measurements were binned and discretized. The bin size, or bin number, is a parameter of the estimation which can introduce bias to the MI calculation. To correct this bias, bin numbers ranging from 5 to 50 were tested and MI plateaued after a bin number of 20. The presented MI was calculated as mean of MIs corresponding to bin numbers of 20-50.

Alternatively, the measured values of mCherry and GFP can be jointly considered as samples of *p(X,Y)* where the input signal is concentration of PhoP inside the cells (approximated via the mCherry measurement) and output signal is the activity of the downstream gene (measured by the value of the reporter GFP). This allows for direct calculation of mutual information between the input and output since marginal probability densities of *X* and *Y* can be calculated from *p(X,Y)*. In this case, mutual information can be calculated from the following equation:

$$I(X;Y) = H(X) + H(Y) - H(Y,X)$$

To calculate the information entropy terms in the above equation, one needs to approximate *p(X)*, *p(Y)*, and *p(X,Y)* based on the paired samples—(mCherry,GFP) measurements. While these probability densities can be approximated through binning similar to calculation of channel capacity described above, we found that the bin size introduces significant bias despite applying bias correction methods such as Miller-Madow (Paninski, 2003) (Fig. S16). Therefore, we resorted to a non-parametric method known as Kraskov-Stögbauer-Grassberger (KSG) (Kraskov et al., 2004) to approximate joint and marginal probability densities. The KSG method relies on finding the Chebyshev distance to k-nearest neighbors of each paired point in the joint space. The distance is then used to count number of neighbors in marginal spaces to calculate joint and marginal probability densities. If the number of neighbors in *X* and *Y* spaces for a certain point in joint space are represented with $n_x$ and $n_y$, respectively, the KSG method calculates mutual information through the following equation:

$$I(X;Y) = \psi(k) + \psi(n) - \frac{1}{n}\sum_{i=1}^{n}\left[\psi\left(n_{x_i}+1\right)+\psi\left(n_{y_i}+1\right)\right]$$

where $k$ and $n$ stand for the number of nearest neighbors in joint space and sample size, respectively, and $\psi(x)$ is the digamma function. The only parameter in the KSG algorithm is the number of nearest neighbors ($k$) which we found not to significantly affect the value of $I(X;Y)$ (Fig. S17). Univariate and multivariate mutual information calculations for continuous variables were performed using scikit-learn and the NPEET Python libraries, respectively.

***Calculating mutual information from dynamic signals***

Mutual information between two continuous dynamic signals—*e.g.*, mCherry and GFP measurements over time in Fig. 2B—can be calculated using the KSG method mentioned above and as described by others (Selimkhanov et al., 2014). One can consider each mCherry or GFP trajectory as an *r*-dimensional point $\boldsymbol{x} = [x_1, \dots, x_r]$ where r is the number of data points sampled uniformly from the trajectory. Following approximation of the joint and marginal distributions of $\boldsymbol{x}$ (mCherry) and $\boldsymbol{y}$ (GFP), mutual information can be calculated using the KSG approach. We found that after a certain threshold (~20), increasing r did not change mutual information significantly (Fig. S18).

To estimate the information encoded in mCherry and GFP joint time series and their correlation with the input condition and cell survival as presented in Figs. 3C and 4G, we used the method described in Jetka *et al*. (Jetka et al., 2019) called Statistical Learning-based Estimation of Mutual Information (SLEMI). SLEMI calculates mutual information between categorical inputs (*e.g.*, input condition or survival quartile) and multivariate outputs (*e.g.*, mCherry, GFP single or joint time series). Jetka *et al*. proposed using a decoding model (*e.g.*, logistic regression) to classify the multivariate outputs into their respective input classes and to compute mutual information using the classification probabilities. That is:

$$I(X;Y) = H(Y) - H(Y|X) = H(X) - H(X|Y)$$

$$I(X;Y) = \int p(x|y)\,p(y)\log\frac{p(x|y)}{p(x)}\,dx\,dy = \int p(x)p(y|x)\log\frac{p(x|y)}{p(x)}\,dx\,dy$$

Similar to the previous section, we assume that the outputs measured for each input group are samples from *p(y|x)*. However, SLEMI has two additional important assumptions: 1. For large sample numbers per group ($n_i$), the integral over $Y$ can be approximated as an average. 2. The marginal input distribution is either known or can be optimized to calculate channel capacity. Consequently:

$$I(X;Y) = \sum_{i=1}^{n} p(x_i) \frac{1}{n_i} \sum_{k=1}^{n_i} \log \frac{p(x_i|y = y_k)}{p(x_i)}$$

In which $p(x_i|y = y_k)$ can be estimated using logistic regression.

We used the SLEMI R-package developed by Jetka *et al*. to compute mutual information values presented in Fig. 3C and 4G. For calculating mutual information between fluorescence signals over time and the input condition or survival quartile, we assumed uniform probability distributions for inputs.

__Supplementary Figures__

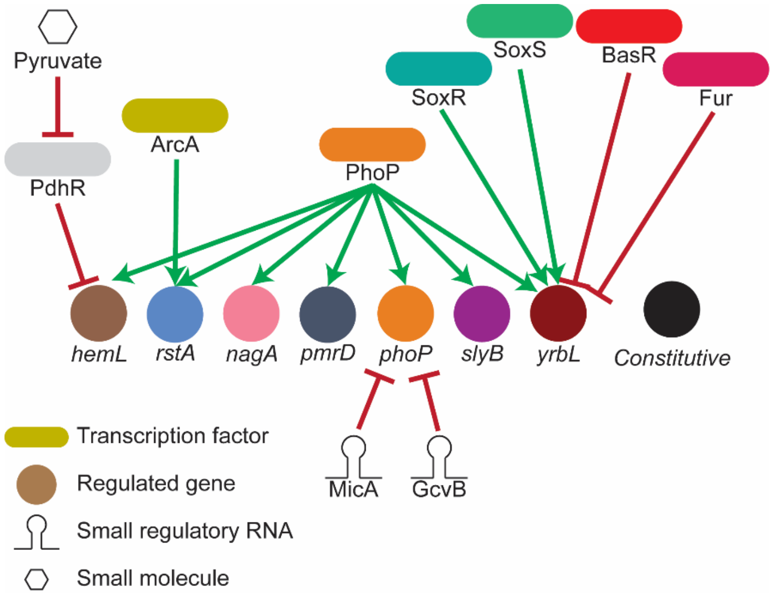


**Figure S1. Schematics of regulators of the PhoP-regulated genes studied in this work.**

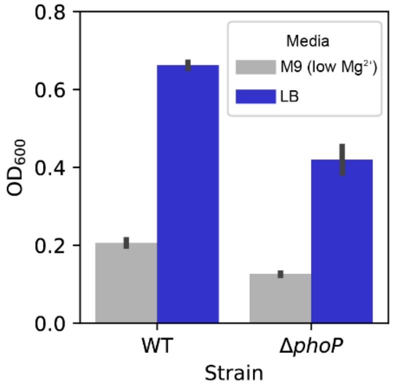


**Figure S2. Comparison of BW25113 (wild-type) and BW25113 Δ*phoP* strains in different media conditions.** Cells were grown in low $Mg^{2+}$ M9 medium (1× M9 salts supplemented with 10 μM $MgSO_4$, 0.4% glucose, and 0.2% casamino acids) or LB Broth. $OD_{600}$ was measured after 5 hours of growth. Error bars indicate standard deviation from n = 2 biological replicates.

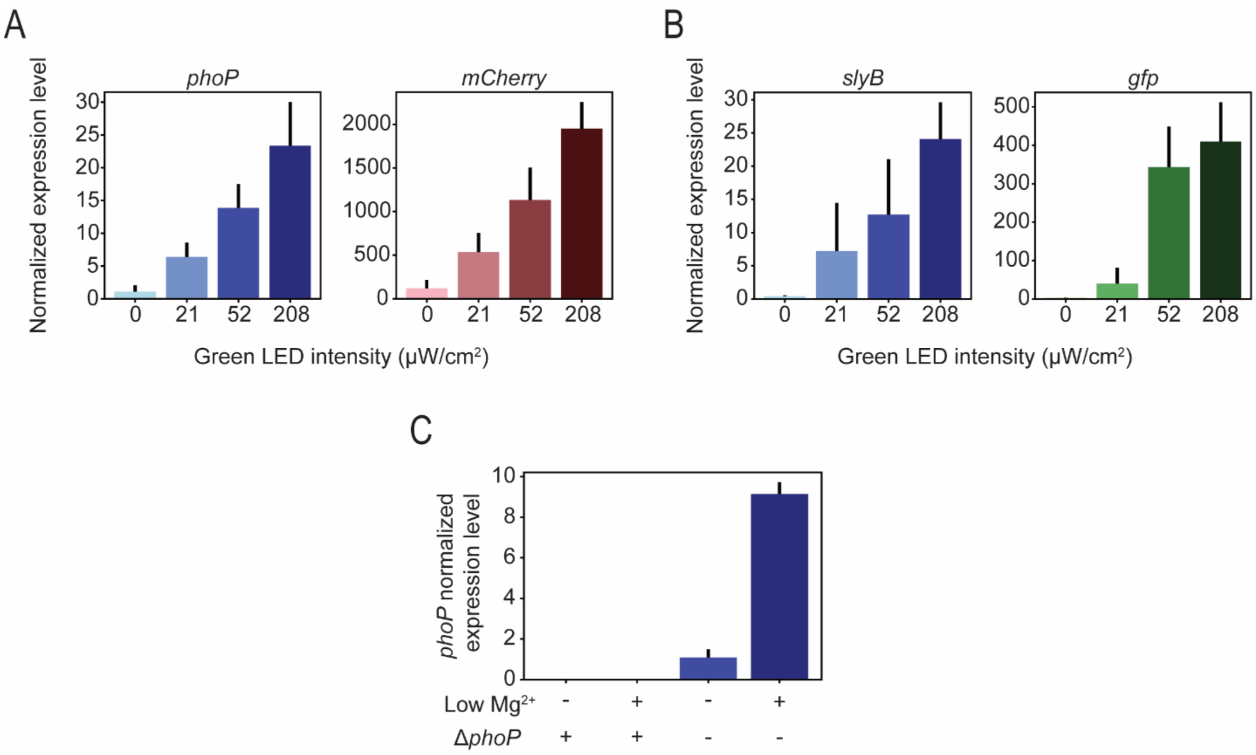


**Figure S3**. **Quantitative PCR confirms the correlation between the abundance of signaling molecules and their fluorescent reporters.** (A) qPCR measurements of *phoP* and *mCherry* relative expression extracted from cells grown while exposed to four different levels of optogenetic stimulation. (B) qPCR measurements of *slyB* and *gfp* relative expression extracted from cells grown while exposed to four different levels of optogenetic stimulation. (C) qPCR measurements of physiological *phoP* relative expression level in wild-type BW25113 cells grown in normal or low $Mg^{2+}$ condition. Also shown is the *phoP* abundance in Δ*phoP* BW25113 cells in both normal and low $Mg2^{+}$ conditions. Data is shown as mean ± standard deviation. n = 3 biological replicates.

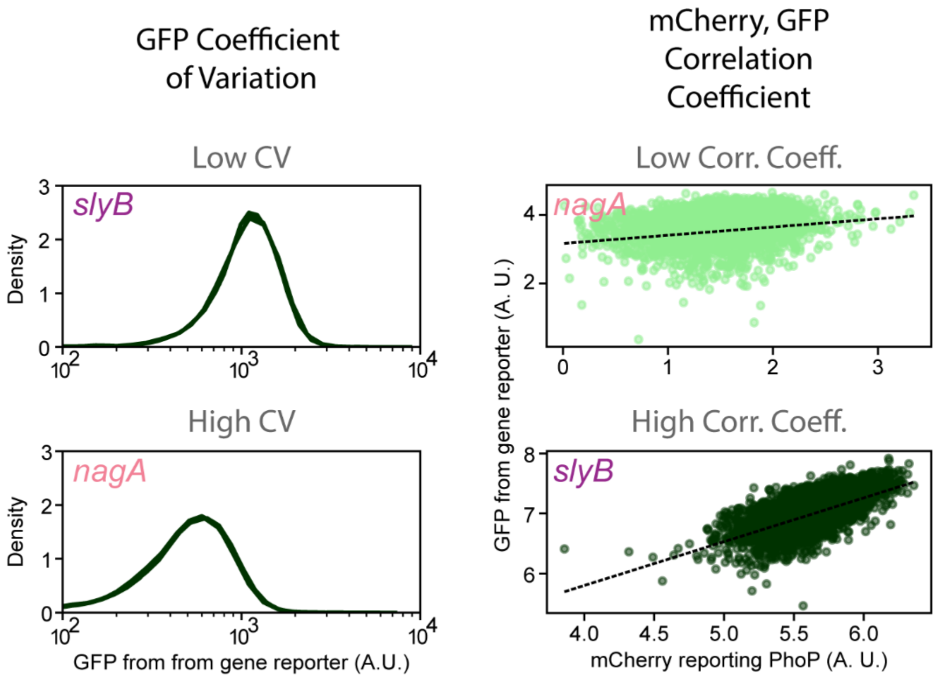


**Figure S4. Representative examples for metrics used for flow cytometry analysis.** Populations with similar mean GFP values can have differences in distribution width (left), which can be quantified by the population coefficient of variation (Fig. 1E). Examples of low coefficient of variation (*slyB,* CV = 0.066 as measured at high stimulation level) and high coefficient of variation (*nagA*, CV = 0.192 as measured at a high stimulation level) are shown. Populations may also have different strengths of correlation between fluorophores regardless of the overall trendline (right), which is quantified by the correlation coefficient (Fig. 1F). Examples of low correlation coefficient (*nagA,* correlation coefficient = 0.21 as measured at low stimulation level) and high correlation coefficient (*slyB*, correlation coefficient = 0.60 as measured at a high stimulation level) are shown. Low and high stimulation levels correspond to 208 μW/cm$^2$ and 0 μW/cm$^2$ green LED intensity, respectively.

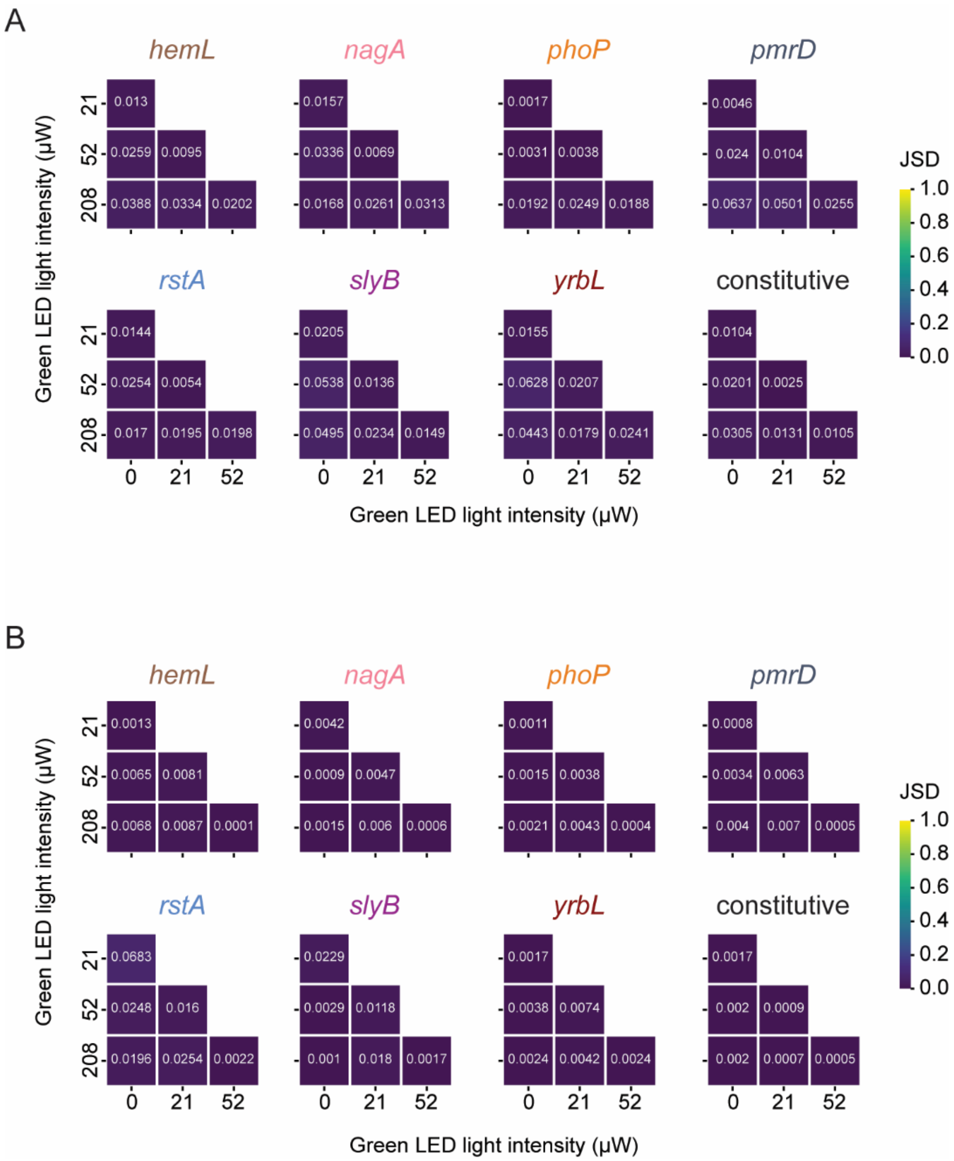


**Figure S5. Pairwise Jensen-Shannon divergence for mCherry and GFP populations shows high distribution similarity.** Pairwise Jensen-Shannon divergence (JSD) between (A) mCherry and (B) GFP populations corresponding to different green LED light intensities per gene. Note that all heatmaps display blue color, meaning JSD values close to zero, indicating high distribution similarity.

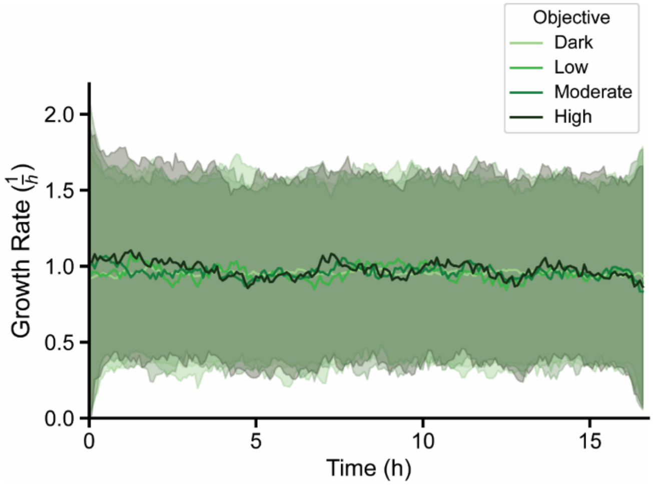


**Figure S6. Impact of prolonged light exposure on cell growth rate.** Growth rate data are shown for cells subjected to either phase contrast imaging alone (Dark) or a combination with phase contrast, GFP, RFP imaging, and DMD illumination consistent with the optogenetic stimulation objectives shown in Fig. 2B.

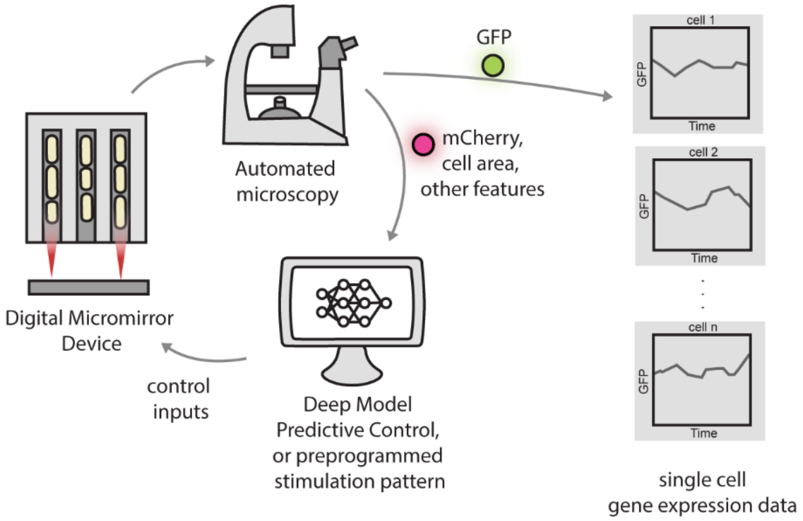


**Figure S7. Schematic of experimental platform.** Cells are cultured in the mother machine microfluidic device and imaged with time-lapse fluorescence microscopy. Fluorescence data and other features such as cell area are extracted on-the-fly and fed into a deep model predictive control algorithm to determine the optimal stimulation pattern for each individual chamber to reach the desired objective. This pattern, along with any preprogrammed stimulation patterns, are enacted on the cells. The cycle then repeats every 5 minutes. GFP data are not used as a control parameter, but are analyzed at the conclusion of the experiment.

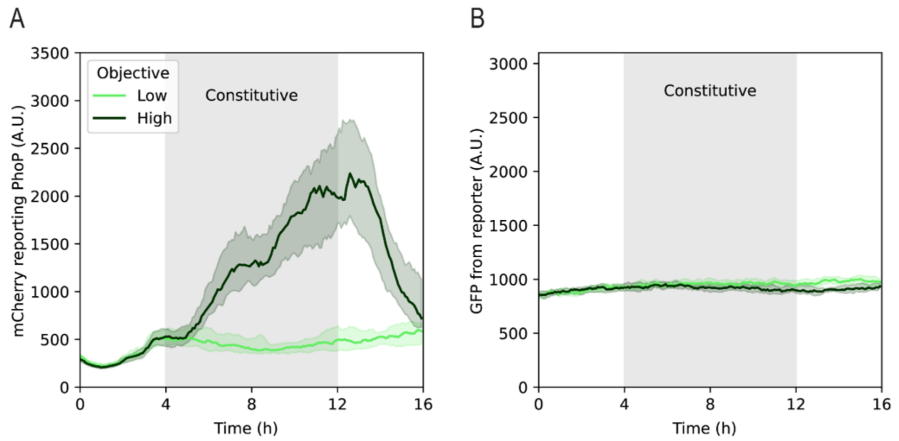


**Figure S8. Single cell control experiments with a constitutive promoter reporter.** (A) mCherry and (B) GFP levels for high and low stimulation regimens.

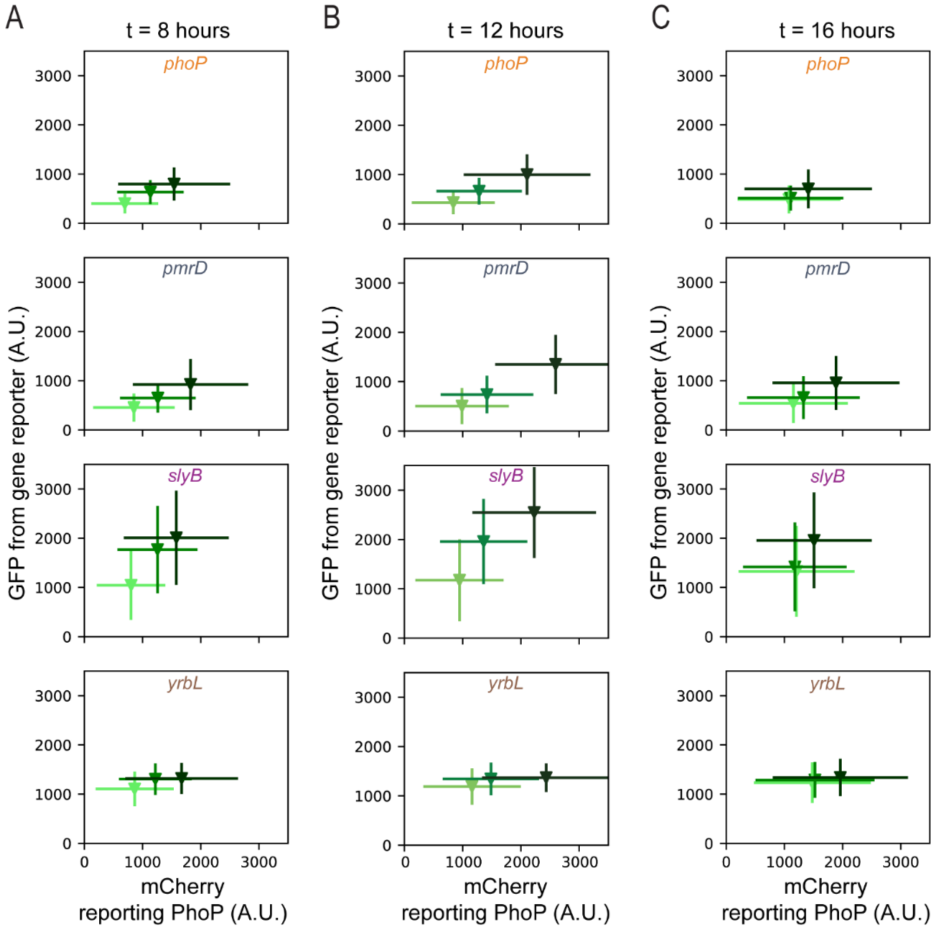


**Figure S9.** Relationship between mean mCherry and mean GFP in all cells at (A) time t = 8, (B) t = 12 hours, and (C) t = 16 hours. n > 300 cells for each reporter (> 100 cells for each stimulation level). Error bars represent standard deviation.

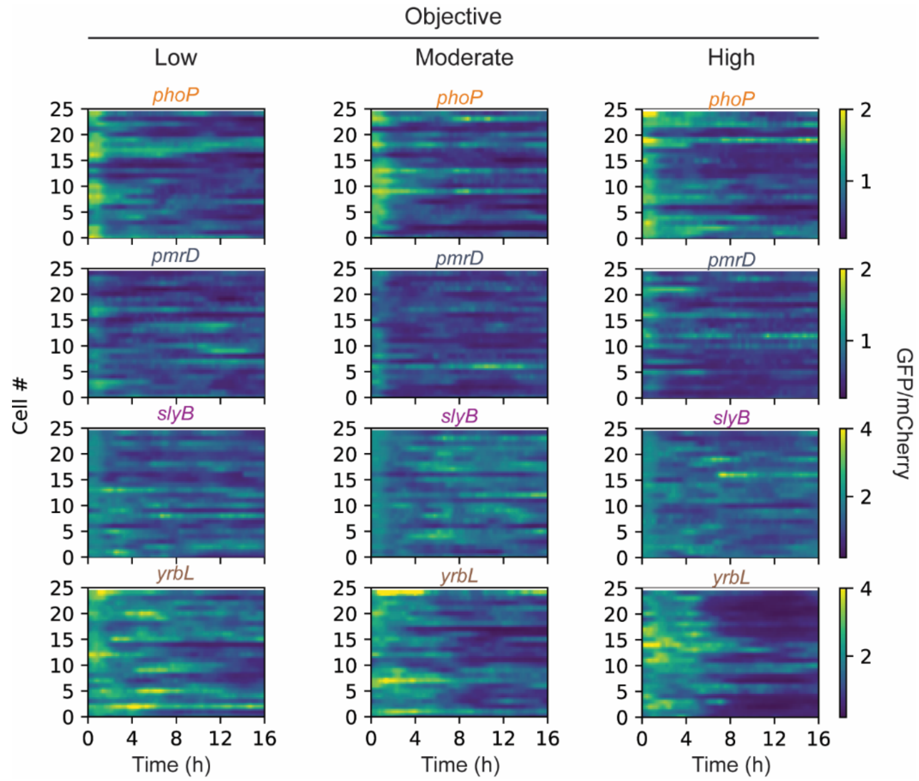


**Figure S10. Temporal and cell-to-cell heterogeneity of joint space over time.** For 25 randomly selected cells from each objective group for each gene, the kymograph shows the ratio of GFP to mCherry over time.

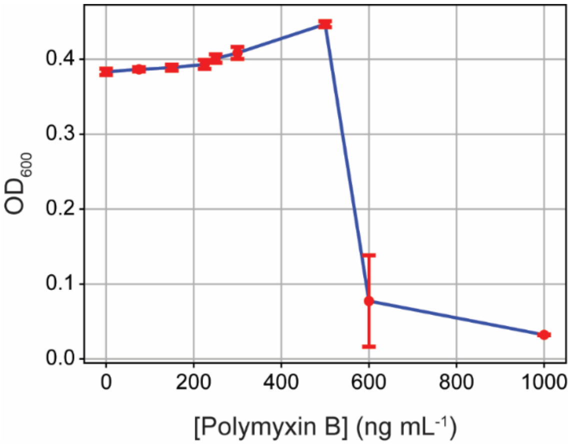


**Figure S11. Minimum inhibitory concentration (MIC) assay verifies lethal concentration of Polymyxin B.** Polymyxin B MIC assay indicating optical density ($A_{600}$) of cells grown in different concentrations of Polymyxin B. Data are shown as mean ± standard deviation from n = 3 biological replicates.

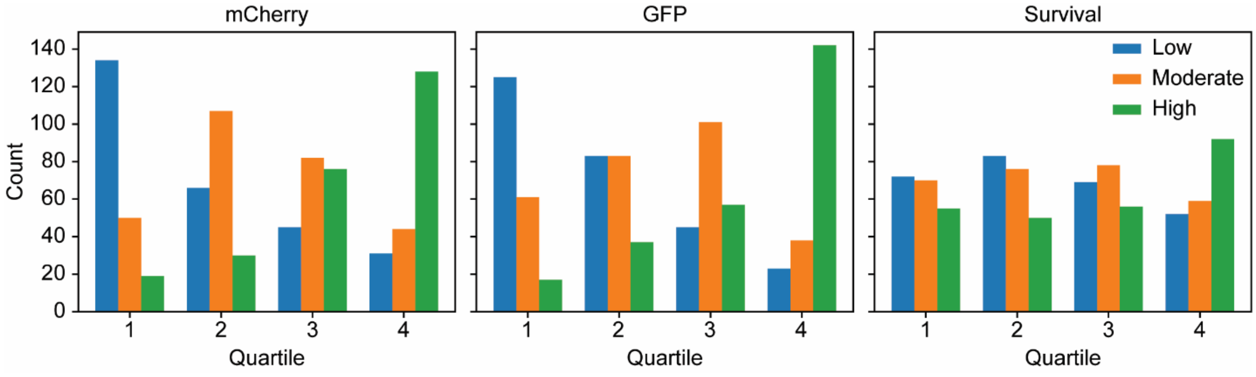


**Figure S12. Segregation of cell mCherry, GFP, and survival data based on optogenetic stimulation group reveals diversity of surviving cells.** Histograms of mCherry (left), GFP (middle), and survival (right) quartiles grouped by optogenetic control objective.

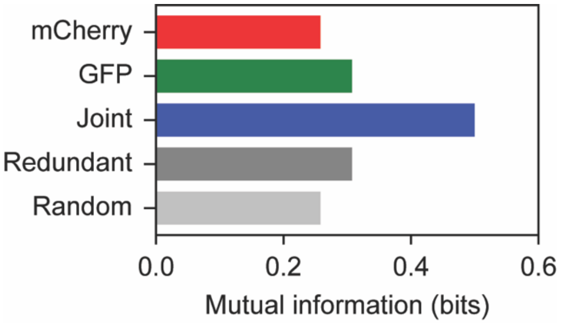


**Figure S13. Addition of redundant or random signals to create joint time series does not increase mutual information.** Mutual information between survival quartile and mCherry time series, GFP time series, or joint time series of concatenated mCherry and GFP (joint), GFP and GFP (redundant), and randomized mCherry and GFP (random).

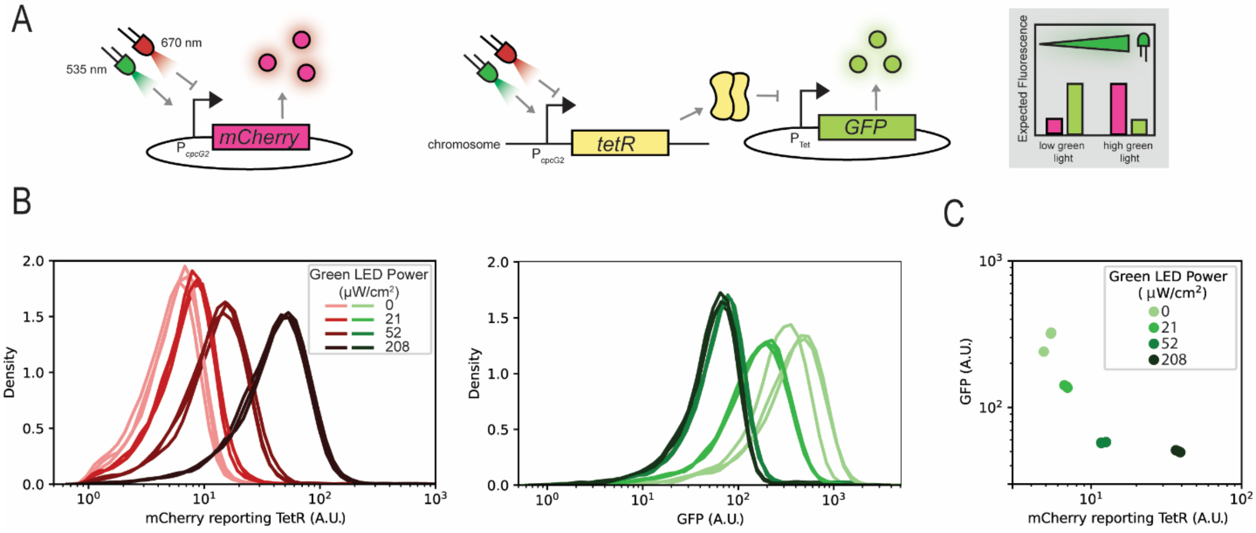


**Figure S14. Optogenetic control of transcription factor dynamics using a synthetic inverter system.** (A) The TetR transcription factor is co-expressed with the fluorophore mCherry under control of the optogenetically-regulated $P_{cpcG2}$ promoter. Red light represses transcription from the promoter, while increasing levels of green light activate higher levels of transcription. A $P_{Tet}$-*gfp* reporter indicates downstream activity of the transcription factor. Also included is a reporter plasmid with the $P_{Tet}$ promoter. With increasing levels of green light, cells will move from a low mCherry, high GFP state to a high mCherry, low GFP state. n = 3 biological replicates for each green light level. (B) mCherry and GFP fluorescence distributions for cells expressing the inverter circuit, under a range of green light stimulation levels. (C) Geometric means of GFP and mCherry fluorescence of the cells from (B).

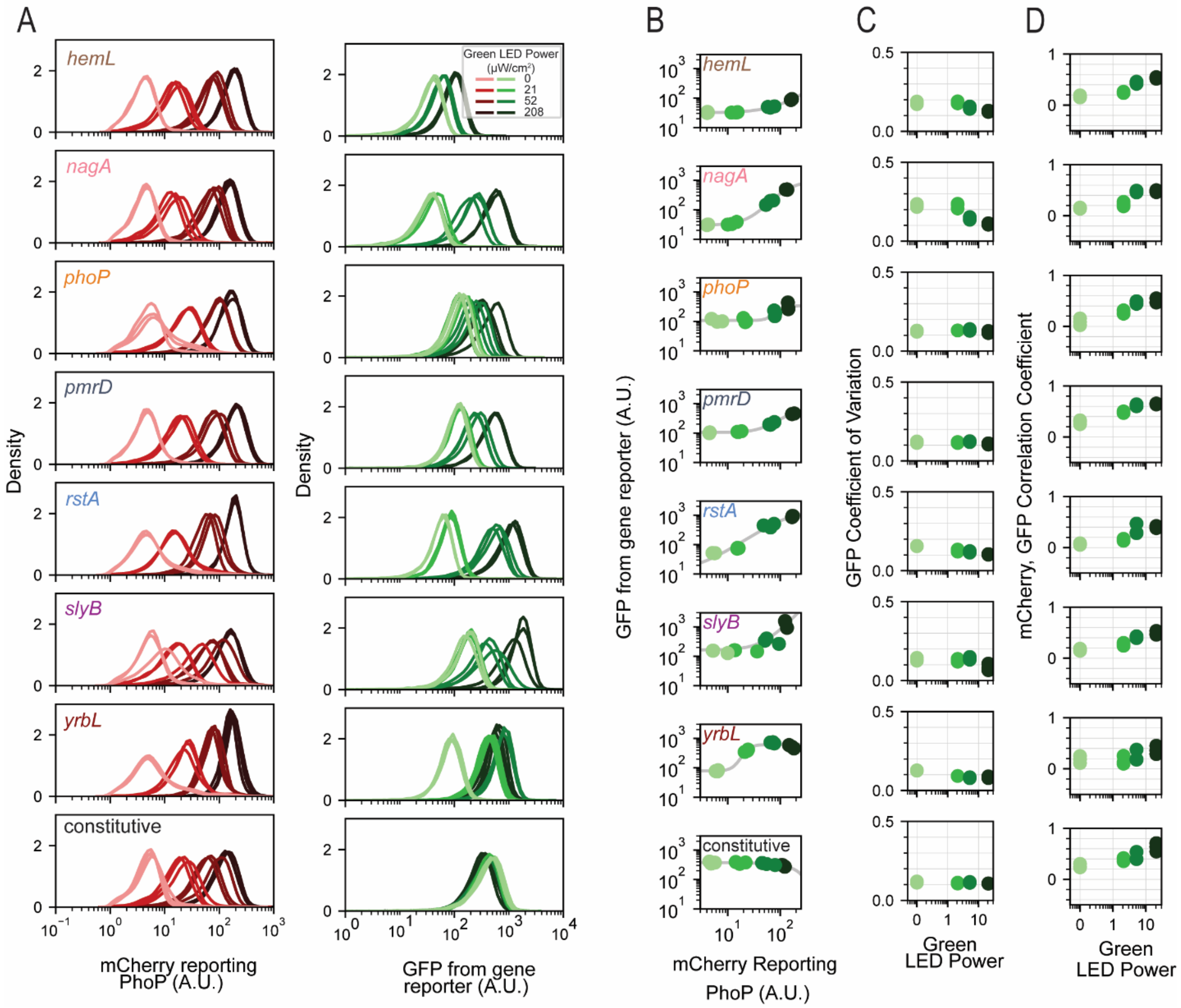


**Figure S15.** (A) mCherry and GFP fluorescence distributions for cells containing reporters for all downstream genes, under a range of green light stimulation levels. n = 3 biological replicates for each green light level. (B) Geometric means of GFP and mCherry fluorescence of the cells from panel A. Grey line represents best fit to a Hill function model. (C) Coefficient of variation for GFP across all cells. (D) Mean correlation coefficient between mCherry and GFP calculated across all individual cells.

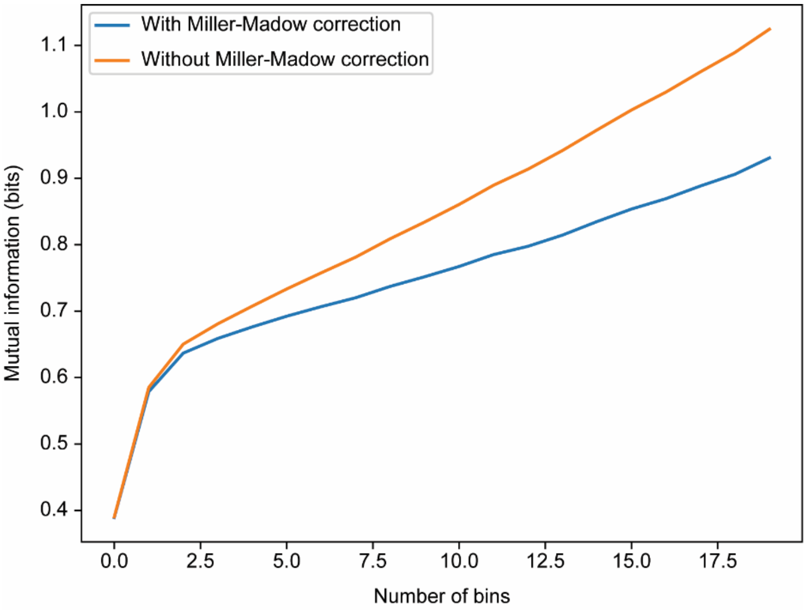


**Figure S16. Calculated mutual information through estimation of joint and marginal probability distributions via binning is influenced by bin number bias.** Plots of mutual information between mCherry (PhoP) and GFP (*phoP*) as a function of number of bins used to discretize the fluorescence populations with or without Miller-Madow correction.

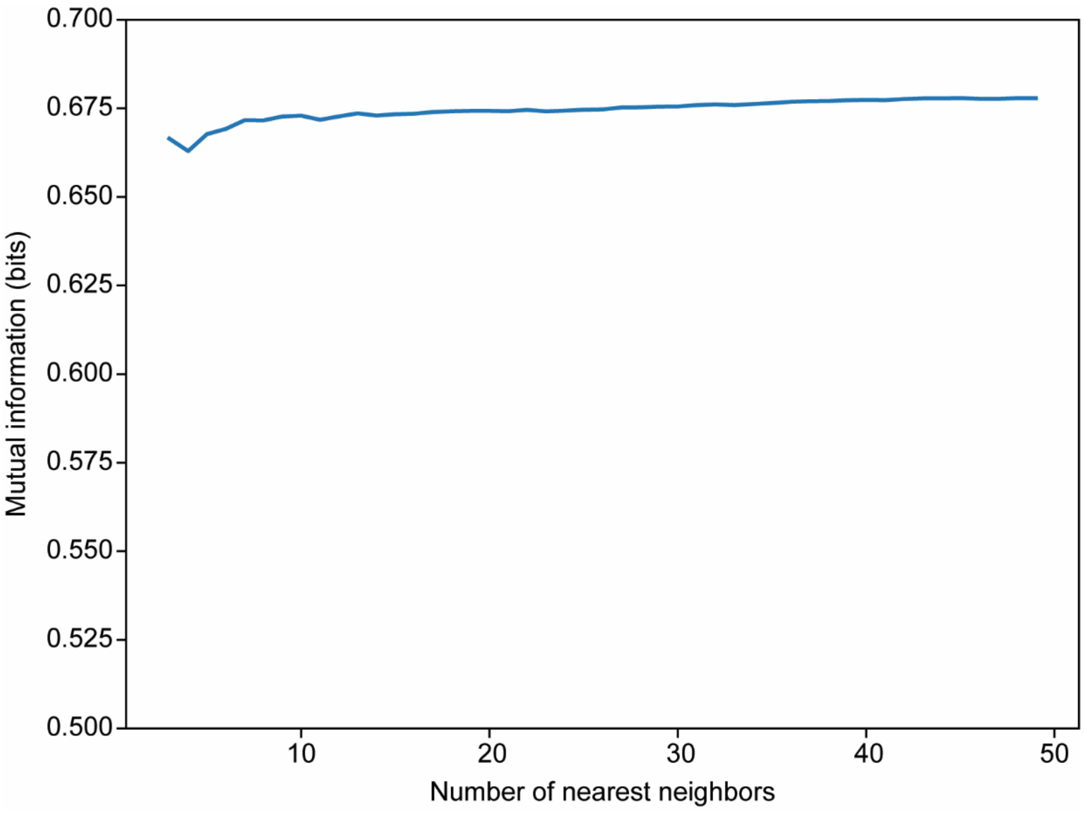


**Figure S17. Kraskov-Stögbauer-Grassberger approach is robust to changes in number of nearest neighbors (k).** Mutual information between mCherry and GFP populations for *pmrD* exhibits only a small change for different *k* values.

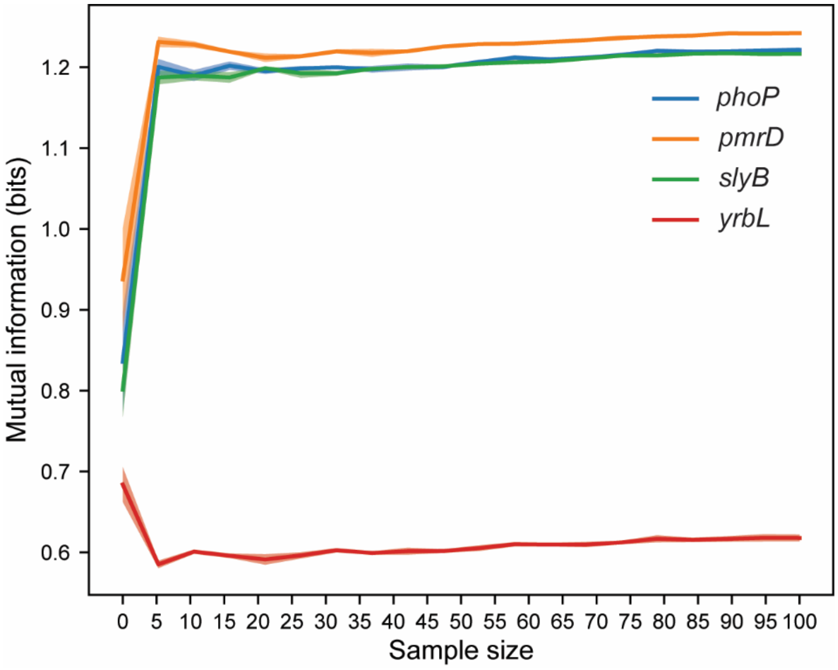


**Figure S18. Mutual information between continuous multivariate mCherry and GFP signals reaches plateau upon sampling only a few points.** Mutual information between dynamic mCherry and GFP signals for different genes reaches a plateau after uniformly sampling the time series with a minimum sample size of 20.

### Supplementary Tables

#### Table S1. Values for Hill equation fit.

| **Reporter** | **A** | **n** | **k** | **B** |
|---|---|---|---|---|
| *hemL* | 52.40 | 3.35 | 37.39 | 30.34 |
| *nagA* | 577.25 | 2.41 | 83.45 | 34.65 |
| *phoP* | 1320.2 | 1.65 | 226.26 | 119.8 |
| *pmrD* | 310.44 | 3.43 | 93.30 | 112.99 |
| *rstA* | 3176.00 | 3.24 | 62.00 | 76.06 |
| *slyB* | 1067.62 | 2.44 | 108.33 | 87.48 |
| *yrbL* | 735.81 | 3.34 | 10.28 | 70.23 |

**Table S2. *p*-values of statistical significance test using bootstrap resampling between coefficients of variation (CV) of green LED intensity groups.** Note that *p*-values are not significant for any of the comparison groups.

| Gene | Group 1 green LED intensity | Group 2 green LED intensity | Group 1 CV | Group 2 CV | *p*-value |
|---|---|---|---|---|---|
| *hemL* | 0 μW/cm$^2$ | 21 μW/cm$^2$ | 0.142 | 0.161 | 0.499 |
| *hemL* | 0 μW/cm$^2$ | 52 μW/cm$^2$ | 0.142 | 0.216 | 0.502 |
| *hemL* | 0 μW/cm$^2$ | 208 μW/cm$^2$ | 0.142 | 0.216 | 0.499 |
| *hemL* | 21 μW/cm$^2$ | 52 μW/cm$^2$ | 0.161 | 0.216 | 0.511 |
| *hemL* | 21 μW/cm$^2$ | 208 μW/cm$^2$ | 0.161 | 0.216 | 0.508 |
| *hemL* | 52 μW/cm$^2$ | 208 μW/cm$^2$ | 0.216 | 0.216 | 0.498 |
| *nagA* | 0 μW/cm$^2$ | 21 μW/cm$^2$ | 0.111 | 0.141 | 0.496 |
| *nagA* | 0 μW/cm$^2$ | 52 μW/cm$^2$ | 0.111 | 0.174 | 0.504 |
| *nagA* | 0 μW/cm$^2$ | 208 μW/cm$^2$ | 0.111 | 0.192 | 0.492 |
| *nagA* | 21 μW/cm$^2$ | 52 μW/cm$^2$ | 0.141 | 0.174 | 0.491 |
| *nagA* | 21 μW/cm$^2$ | 208 μW/cm$^2$ | 0.141 | 0.192 | 0.498 |
| *nagA* | 52 μW/cm$^2$ | 208 μW/cm$^2$ | 0.174 | 0.192 | 0.498 |
| *phoP* | 0 μW/cm$^2$ | 21 μW/cm$^2$ | 0.110 | 0.139 | 0.506 |
| *phoP* | 0 μW/cm$^2$ | 52 μW/cm$^2$ | 0.110 | 0.134 | 0.505 |
| *phoP* | 0 μW/cm$^2$ | 208 μW/cm$^2$ | 0.110 | 0.134 | 0.495 |
| *phoP* | 21 μW/cm$^2$ | 52 μW/cm$^2$ | 0.139 | 0.134 | 0.502 |
| *phoP* | 21 μW/cm$^2$ | 208 μW/cm$^2$ | 0.139 | 0.134 | 0.495 |
| *phoP* | 52 μW/cm$^2$ | 208 μW/cm$^2$ | 0.134 | 0.134 | 0.506 |
| *pmrD* | 0 μW/cm$^2$ | 21 μW/cm$^2$ | 0.124 | 0.138 | 0.488 |
| *pmrD* | 0 μW/cm$^2$ | 52 μW/cm$^2$ | 0.124 | 0.134 | 0.491 |
| *pmrD* | 0 μW/cm$^2$ | 208 μW/cm$^2$ | 0.124 | 0.136 | 0.505 |
| *pmrD* | 21 μW/cm$^2$ | 52 μW/cm$^2$ | 0.138 | 0.134 | 0.497 |
| *pmrD* | 21 μW/cm$^2$ | 208 μW/cm$^2$ | 0.138 | 0.136 | 0.504 |
| *pmrD* | 52 μW/cm$^2$ | 208 μW/cm$^2$ | 0.134 | 0.136 | 0.506 |
| *rstA* | 0 μW/cm$^2$ | 21 μW/cm$^2$ | 0.058 | 0.126 | 0.504 |
| *rstA* | 0 μW/cm$^2$ | 52 μW/cm$^2$ | 0.058 | 0.146 | 0.509 |
| *rstA* | 0 μW/cm$^2$ | 208 μW/cm$^2$ | 0.058 | 0.162 | 0.496 |
| *rstA* | 21 μW/cm$^2$ | 52 μW/cm$^2$ | 0.126 | 0.146 | 0.506 |
| *rstA* | 21 μW/cm$^2$ | 208 μW/cm$^2$ | 0.126 | 0.162 | 0.499 |
| *rstA* | 52 μW/cm$^2$ | 208 μW/cm$^2$ | 0.146 | 0.162 | 0.500 |
| *slyB* | 0 μW/cm$^2$ | 21 μW/cm$^2$ | 0.066 | 0.101 | 0.503 |
| *slyB* | 0 μW/cm$^2$ | 52 μW/cm$^2$ | 0.066 | 0.106 | 0.495 |
| *slyB* | 0 μW/cm$^2$ | 208 μW/cm$^2$ | 0.066 | 0.109 | 0.501 |
| *slyB* | 21 μW/cm$^2$ | 52 μW/cm$^2$ | 0.101 | 0.106 | 0.500 |
| *slyB* | 21 μW/cm$^2$ | 208 μW/cm$^2$ | 0.101 | 0.109 | 0.498 |
| *slyB* | 52 μW/cm$^2$ | 208 μW/cm$^2$ | 0.106 | 0.109 | 0.503 |
| *yrbL* | 0 μW/cm$^2$ | 21 μW/cm$^2$ | 0.083 | 0.069 | 0.499 |
| *yrbL* | 0 μW/cm$^2$ | 52 μW/cm$^2$ | 0.083 | 0.093 | 0.497 |
| *yrbL* | 0 μW/cm$^2$ | 208 μW/cm$^2$ | 0.083 | 0.119 | 0.499 |
| *yrbL* | 21 μW/cm$^2$ | 52 μW/cm$^2$ | 0.069 | 0.093 | 0.506 |
| *yrbL* | 21 μW/cm$^2$ | 208 μW/cm$^2$ | 0.069 | 0.119 | 0.505 |
| *yrbL* | 52 μW/cm$^2$ | 208 μW/cm$^2$ | 0.093 | 0.119 | 0.496 |
| Constitutive | 0 μW/cm$^2$ | 21 μW/cm$^2$ | 0.079 | 0.078 | 0.497 |
| Constitutive | 0 μW/cm$^2$ | 52 μW/cm$^2$ | 0.079 | 0.079 | 0.493 |
| Constitutive | 0 μW/cm$^2$ | 208 μW/cm$^2$ | 0.079 | 0.079 | 0.504 |
| Constitutive | 21 μW/cm$^2$ | 52 μW/cm$^2$ | 0.078 | 0.079 | 0.498 |
| Constitutive | 21 μW/cm$^2$ | 208 μW/cm$^2$ | 0.078 | 0.079 | 0.506 |
| Constitutive | 52 μW/cm$^2$ | 208 μW/cm$^2$ | 0.079 | 0.079 | 0.498 |

**Table S3. *p*-values of statistical significance test using z-test between Fisher transformed mCherry-GFP correlation coefficients (*r*) of green LED intensity groups.**

| Gene | Group 1 green LED intensity | Group 2 green LED intensity | Group 1 *r* | Group 2 *r* | *p*-value |
|---|---|---|---|---|---|
| *hemL* | 0 µW/cm$^2$ | 21 µW/cm$^2$ | 0.283 | 0.37 | 1.38E-14 |
| *hemL* | 0 µW/cm$^2$ | 52 µW/cm$^2$ | 0.283 | 0.27 | 0.3155 |
| *hemL* | 0 µW/cm$^2$ | 208 µW/cm$^2$ | 0.283 | 0.257 | 0.0372 |
| *hemL* | 21 µW/cm$^2$ | 52 µW/cm$^2$ | 0.37 | 0.27 | 0 |
| *hemL* | 21 µW/cm$^2$ | 208 µW/cm$^2$ | 0.37 | 0.257 | 0 |
| *hemL* | 52 µW/cm$^2$ | 208 µW/cm$^2$ | 0.27 | 0.257 | 0.2666 |
| *nagA* | 0 µW/cm$^2$ | 21 µW/cm$^2$ | 0.518 | 0.518 | 0.9906 |
| *nagA* | 0 µW/cm$^2$ | 52 µW/cm$^2$ | 0.518 | 0.365 | 0 |
| *nagA* | 0 µW/cm$^2$ | 208 µW/cm$^2$ | 0.518 | 0.214 | 0 |
| *nagA* | 21 µW/cm$^2$ | 52 µW/cm$^2$ | 0.518 | 0.365 | 0 |
| *nagA* | 21 µW/cm$^2$ | 208 µW/cm$^2$ | 0.518 | 0.214 | 0 |
| *nagA* | 52 µW/cm$^2$ | 208 µW/cm$^2$ | 0.365 | 0.214 | 0 |
| *phoP* | 0 µW/cm$^2$ | 21 µW/cm$^2$ | 0.737 | 0.615 | 0 |
| *phoP* | 0 µW/cm$^2$ | 52 µW/cm$^2$ | 0.737 | 0.552 | 0 |
| *phoP* | 0 µW/cm$^2$ | 208 µW/cm$^2$ | 0.737 | 0.382 | 0 |
| *phoP* | 21 µW/cm$^2$ | 52 µW/cm$^2$ | 0.615 | 0.552 | 1.71E-13 |
| *phoP* | 21 µW/cm$^2$ | 208 µW/cm$^2$ | 0.615 | 0.382 | 0 |
| *phoP* | 52 µW/cm$^2$ | 208 µW/cm$^2$ | 0.552 | 0.382 | 0 |
| *pmrD* | 0 µW/cm$^2$ | 21 µW/cm$^2$ | 0.562 | 0.527 | 1.43E-04 |
| *pmrD* | 0 µW/cm$^2$ | 52 µW/cm$^2$ | 0.562 | 0.459 | 0 |
| *pmrD* | 0 µW/cm$^2$ | 208 µW/cm$^2$ | 0.562 | 0.206 | 0 |
| *pmrD* | 21 µW/cm$^2$ | 52 µW/cm$^2$ | 0.527 | 0.459 | 4.19E-12 |
| *pmrD* | 21 µW/cm$^2$ | 208 µW/cm$^2$ | 0.527 | 0.206 | 0 |
| *pmrD* | 52 µW/cm$^2$ | 208 µW/cm$^2$ | 0.459 | 0.206 | 0 |
| *rstA* | 0 µW/cm$^2$ | 21 µW/cm$^2$ | 0.522 | 0.496 | 0.0061 |
| *rstA* | 0 µW/cm$^2$ | 52 µW/cm$^2$ | 0.522 | 0.442 | 2.22E-16 |
| *rstA* | 0 µW/cm$^2$ | 208 µW/cm$^2$ | 0.522 | 0.279 | 0 |
| *rstA* | 21 µW/cm$^2$ | 52 µW/cm$^2$ | 0.496 | 0.442 | 5.99E-08 |
| *rstA* | 21 µW/cm$^2$ | 208 µW/cm$^2$ | 0.496 | 0.279 | 0 |
| *rstA* | 52 µW/cm$^2$ | 208 µW/cm$^2$ | 0.442 | 0.279 | 0 |
| *slyB* | 0 µW/cm$^2$ | 21 µW/cm$^2$ | 0.605 | 0.572 | 7.31E-05 |
| *slyB* | 0 µW/cm$^2$ | 52 µW/cm$^2$ | 0.605 | 0.427 | 0 |
| *slyB* | 0 µW/cm$^2$ | 208 µW/cm$^2$ | 0.605 | 0.194 | 0 |
| *slyB* | 21 µW/cm$^2$ | 52 µW/cm$^2$ | 0.572 | 0.427 | 0 |
| *slyB* | 21 µW/cm$^2$ | 208 µW/cm$^2$ | 0.572 | 0.194 | 0 |
| *slyB* | 52 µW/cm$^2$ | 208 µW/cm$^2$ | 0.427 | 0.194 | 0 |
| *yrbL* | 0 µW/cm$^2$ | 21 µW/cm$^2$ | 0.468 | 0.363 | 0 |
| *yrbL* | 0 µW/cm$^2$ | 52 µW/cm$^2$ | 0.468 | 0.364 | 0 |
| *yrbL* | 0 µW/cm$^2$ | 208 µW/cm$^2$ | 0.468 | 0.279 | 0 |
| *yrbL* | 21 µW/cm$^2$ | 52 µW/cm$^2$ | 0.363 | 0.364 | 0.8932 |
| *yrbL* | 21 µW/cm$^2$ | 208 µW/cm$^2$ | 0.363 | 0.279 | 3.86E-13 |
| *yrbL* | 52 µW/cm$^2$ | 208 µW/cm$^2$ | 0.364 | 0.279 | 2.43E-13 |
| Constitutive | 0 µW/cm$^2$ | 21 µW/cm$^2$ | 0.714 | 0.626 | 0 |
| Constitutive | 0 µW/cm$^2$ | 52 µW/cm$^2$ | 0.714 | 0.586 | 0 |
| Constitutive | 0 µW/cm$^2$ | 208 µW/cm$^2$ | 0.714 | 0.464 | 0 |
| Constitutive | 21 µW/cm$^2$ | 52 µW/cm$^2$ | 0.626 | 0.586 | 2.96E-06 |
| Constitutive | 21 µW/cm$^2$ | 208 µW/cm$^2$ | 0.626 | 0.464 | 0 |
| Constitutive | 52 µW/cm$^2$ | 208 µW/cm$^2$ | 0.586 | 0.464 | 0 |

**Table S4. List of qPCR primers for target genes.**

| Target gene | Primer sequence | |
|---|---|---|
| *phoP* | FWD | CGATGTTTCACTGCCGATTCTGG |
| | REV | CGCATTAATGCCTGCATTCGC |
| *mCherry* | FWD | CGTCTAGCGAACGCATGTATCC |
| | REV | TAATATTCACATTGTACGCGCCAGG |
| *slyB* | FWD | GGCACCATCGTTAACGTACGT |
| | REV | ATTGCACTCTGTACGCCCTGA |
| *gfp* | FWD | CTACAAGACACGTGCTGAAGTCAAG |
| | REV | GAACGCTTCCATCTTCAATGTTGTG |
| *gapA* | FWD | GGGCTACACCGAAGATGACGTA |
| | REV | GGAGTAACCGGTTTCGTTGTCG |
